# Actin-based protrusions lead microtubules during stereotyped axon initiation in spinal neurons *in vivo*

**DOI:** 10.1101/2020.10.24.353284

**Authors:** Rachel E. Moore, Sînziana Pop, Caché Alleyne, Jonathan DW Clarke

## Abstract

*In vitro*, developing neurons progress through well-defined stages to form an axon and multiple dendrites. *In vivo*, neurons are derived from progenitors within a polarised neuroepithelium and it is not clear how axon initiation observed *in vitro* relates to what occurs in a complex, three-dimensional *in vivo* environment. Here we show that the position of axon initiation in embryonic zebrafish spinal neurons is extremely consistent across neuronal sub-types. We investigated what mechanisms may regulate axon positioning *in vivo* and found that microtubule organising centres are located distant from the site of axon initiation in contrast to that observed *in vitro*, and that microtubule plus-ends are not enriched in the axon during axon initiation. F-actin accumulation precedes axon formation and nascent axons form but are not stabilised in the absence of microtubules. Laminin depletion removes a spatial cue for axon initiation but axon initiation remains robust.

## INTRODUCTION

During development neurons polarise to form axonal and dendritic domains. This is essential for circuit formation and for the directional movement of information through the nervous system, but many of the fundamental mechanisms that initiate and build axons and dendrites are not understood.

*In vitro*, dissociated rodent hippocampal neurons progress through well-defined stages to develop one axon and multiple dendrites ((Dotti et al., 1988); reviewed in). The neuron first extends several neurites, any of which has the potential to become an axon. Neuronal polarisation occurs when one neurite is specified to become the axon and grows longer and faster than the others, which subsequently become dendrites.

The question of which neurite becomes the axon has been studied extensively. Proteins associated with axon specification (also called axon determinants, including PIP3, LKB1 and the Par complex) move between neurites before becoming persistently localised in one neurite, which subsequently becomes the axon (Shi et al., 2003; Ménager et al., 2004; Yoshimura et al., 2006; Barnes et al., 2007; Shelly et al., 2007). Their downstream signalling pathways reinforce axonal identity and induce cytoskeletal changes. Neuronal polarisation can occur stochastically in the absence of extracellular factors, but environmental cues such as growth and guidance factors (Hilliard and Bargmann, 2006; Wolman et al., 2008; Yi et al., 2010; Cheng et al., 2011; Nakamuta et al., 2011; Shelly et al., 2011) and extracellular matrix proteins (Gomez and Letourneau, 1994; Esch et al., 1999; Ménager et al., 2004; Randlett et al., 2011) can position axogenesis by regulating the localisation of axon determinants. However, one caveat of using this *in vitro* system is that axon initiation is defined by the growth and specification of an already existing neurite. Indeed, axon growth is required to identify neuronal polarisation and axon initiation in this context (Goslin and Banker, 1989; Yamamoto et al., 2012). As such, it is difficult to determine what is actually required for axonal specification and what simply promotes axonal growth (reviewed in (Barnes and Polleux, 2009)).

*In vitro*, an increase in intracellular trafficking towards the future axon early in neuronal polarisation is thought to be due to polarisation of the microtubule cytoskeleton and to cause the accumulation of axon determinants at the nascent axon (Bradke and Dotti, 1997). Polarisation of the microtubule array has been suggested to be driven by the asymmetric localisation of microtubule organising centres (MTOCs) - the centrosome and Golgi complex - which nucleate microtubules. Indeed, both the centrosome and Golgi complex have been described to localise to the base of the axon in hippocampal neurons (de Anda et al., 2005), and cerebellar granule cells *in vitro* (Zmuda and Rivas, 1998). However, other studies have described the centrosome as positioned randomly with respect to the axon in cultured hippocampal neurons (Dotti and Banker, 1991), or to localise to the future axon after it has developed (Gärtner et al., 2012). *In vivo*, the centrosome of Rohon-Beard neurons in the embryonic zebrafish spinal cord is on the opposite side of the cell to the central axons when they are initiated, but has been reported to then move to the base of the peripheral axon during its initiation (Andersen and Halloran, 2012). The centrosome is not close to the axon in zebrafish retinal ganglion cells (Zolessi et al., 2006) or tegmental hindbrain neurons (Distel et al., 2010), and neurons in *Drosophila* embryos without centrosomes are able to extend axons apparently normally (Basto et al., 2006).

We have recently shown that the well-defined stages of neuronal polarisation described *in vitro* do not occur in neurons in the embryonic zebrafish neural tube (Hadjivasiliou et al., 2019). The neuronal soma moves to the basal surface of the neuroepithelium while maintaining an attachment to the apical surface and then extends two long, transient protrusions rostrally and caudally along the basal surface of the neural tube. In contrast to the neurites observed in polarising neurons in culture (Dotti et al., 1988), these protrusions have stereotyped orientations, are transient and are fully retracted along with the apical attachment prior to axon extension. The axon is then initiated directly from the cell body rather than developing from a pre-existing neurite, and well before the appearance of dendrites. In the current study we focused on the stage after basal protrusion retraction to investigate what regulates the initiation of axonal outgrowth in the embryonic spinal neurons. We used time-lapse imaging and observed that the axon is consistently initiated from the basal and ventral side of the soma. Laminin promotes the basal orientation of axon initiation but we were not able to perturb the ventral orientation of axon initiation, which is particularly robust. MTOCs are at the opposite side of the cell at the time of axon initiation and F-actin localises to the future axon initiation site before the accumulation of microtubule plus-ends. Nascent axons can form in the absence of microtubules. This study establishes embryonic zebrafish spinal neurons as a favourable model for investigating axon initiation *in vivo* and proposes F-actin accumulation as the primary cytoskeletal element for axon initiation.

## METHODS

Wildtype (WT; AB), transgenic (TgBAC(*Pard3-GFP*) (Symonds et al., 2020); Tg(*cdh2:cdh2-tFT*) (Revenu et al., 2014); Tg(arl13b-GFP) (Borovina et al., 2010); Tg(actb1:*utr-mCherry*) (Krens et al., 2017)) and mutant (*Sly/lamC1* (Kettleborough et al., 2013); *Smu*^*b641*^/*Smoothened* (Barresi et al., 2010)) zebrafish lines were maintained under standard conditions (Westerfield, 2000). Embryos were raised in aquarium water at 28.5°C.

To observe individual cells, we injected zebrafish embryos at 32-64 cell stage with mRNA encoding fluorescently-tagged proteins: EGFP-CAAX (Kwan et al., 2007), mKate-CAAX (Hadjivasiliou et al., 2019), H2B-RFP (Megason and Fraser, 2003), lifeact-Ruby (Riedl et al., 2008), Kif5c560-YFP (Randlett et al., 2011), EB3-GFP (Norden et al., 2009), centrin2-EGFP (Distel et al., 2010), centrin2-RFP, GM130-EGFP or GM130-RFP (Durdu et al., 2014). Occasionally we coinjected mRNA coding for dominant negative Suppressor of Hairless (dnSuH; (Wettstein et al., 1997)) to increase the likelihood of labelled cells differentiating into neurons. WT or Tg(actb1:*utr-mCherry*) embryos were injected at one-cell stage with 3.4 ng of LamininC1 morpholino (lamMO; 5′-TGTGCCTTTTGCTATTGCGACCTC-3′; (Parsons et al., 2002)) to disrupt laminin expression.

For immunohistochemistry, embryos were dechorionated, anaesthetised with MS-222 (Sigma-Aldrich/Merck, St. Louis, U.S.A.) and fixed in 4% PFA overnight at 4°C. Blocking was performed for two hours at room temperature in appropriate serum. Antibodies were diluted in blocking solution. Embryos were incubated in primary antibody overnight at 4° C (chick α-GFP, Abcam, Cat# AB13970, Lot# GF305729-1; mouse α- γ-tubulin, Sigma-Aldrich, Cat# T6557, Lot# 066M4858V; rabbit α- α- tubulin, Abcam, Cat# AB233661) and in secondary antibody for two hours at room temperature (Alexa goat α-chick 488, Life Technologies, Cat# A11039, Lot# 1812246; Alexa goat α-mouse 568, Life Technologies, Cat# A1104, Lot# 1863187; Alexa goat α- rabbit 633, Life Technologies, Cat# A21071, Lot# 558885).

Imaging was performed from 16 hpf. Embryos were dechorionated, mounted in low-melting point agarose (Sigma-Aldrich) and anaesthetised with MS-222 (Sigma-Aldrich/Merck) if required (Alexandre et al., 2010). Confocal imaging was performed on a spinning disk confocal (PerkinElmer, Waltham, U.S.A.) or LSM880 laser scanning confocal (Zeiss, Oberkochen, Germany) with or without Airyscan, using a 20x water immersion objective with numerical aperture of 0.95 or higher. For high resolution imaging we used Zeiss Airyscan acquisition and processing. Lightsheet imaging was performed on a Zeiss LightSheet Z.1 microscope using 10x illumination objectives and 20x water immersion detection objectives. If required, nocodazole (5 mg/mL stock in DMSO) was diluted in fish water to a final concentration of 5 μg/mL. Following treatment, nocodazole was washed out of the imaging chamber with fish water.

Images were acquired as 40-100 um deep z-stacks with or without time-lapse every 2 seconds to 10 minutes over 5-15 hours depending on the experiment. Images and videos in the manuscript result from maximum projections of z-stacks using Fiji (Schindelin et al., 2012) or 3D reconstructions using Volocity (Perkin Elmer). Surrounding cells were occasionally edited from the field of view using ImageJ or Imaris (Bitplane, Belfast, U.K.) to more clearly show behaviours of the individual cells under investigation.

To analyse organelle position with respect to the cell centroid, the field of view was reoriented so the basal surface was to the right. The cell centroid was determined using the ImageJ 3D Object Counter plugin and the 3D coordinates of the site of axon initiation, centrosome or Golgi complex was determined manually. Trigonometry was used to calculate the distance and angle of each organelle with respect to the cell centroid and this was analysed using Moore’s modification of the Rayleigh statistical test. Trigonometry was also used to calculate the distance between two different organelles within the same cell. These positions were analysed using Moore’s test for paired data. Organelle positions in different conditions were compared using Batschelet’s alternative to the Hotelling test.

Distances between organelles in different conditions were analysed using Student’s t-test except for centrosome-axon distance in *Sly*^−/−^ and lamMO-injected embryos, which were compared to WT using one-way ANOVA. Change in cilium length over time was measured using ImageJ and analysed using non-linear regression to compare each slope to 0 (Prism 8, GraphPad, San Diego, U.S.A.). Protrusion length, width and duration were measured using ImageJ and compared using Student’s t-test. Fluorescence intensity analysis was performed using the ImageJ Plot Profile plugin. Distances between neuron and basal surface were compared using the Student’s t-test.

## RESULTS

### Axon initiation *in vivo* is highly stereotyped

To observe neuronal polarisation *in vivo*, we sparsely and randomly labelled zebrafish embryonic spinal cord cells with a membrane marker and imaged them using time-lapse confocal microscopy from 16 hours post fertilisation (hpf). We previously reported that differentiating neurons in the zebrafish spinal cord go through a distinctive and very stereotyped T-shaped morphology before extending an axon (Figure 1B; Video 01; (Hadjivasiliou et al., 2019)). To better understand axon initiation, we focussed on the time immediately after retraction of the basal protrusions and apical attachment (e.g. from 19h53 in Figure 1B). At this initial stage neurons had no prominent or long-lasting protrusions but extended small, transient protrusions and filopodia (Figure 1C −2h to −20m; Video 02). These transient protrusions occurred in many directions at first (e.g. Figure 1C, −1h30) but were gradually restricted to a more defined baso-ventral position on the cell body. Each neuron then extended a single axonal protrusion from this position (Figure 1C, 0h). This protrusion was stable and did not grow significantly for approximately 30 minutes (Figure 1C 0h to 30m; Figure 1 – figure supplement 1) before extending rapidly (Figure 1C 45m to 2h20). We defined the time of axon initiation as the first time point that showed a persistent protrusion that subsequently matured into an axon with growth cone (Figure 1C, 0h). We call this persistent protrusion the nascent axon. We defined the site of axon initiation as the position that the nascent axon protrusion emerges from the cell body at this time point (Figure 1C 0h, asterisk). The position of the nascent axon was analysed with respect to the cell centroid and found to be highly biased towards the baso-ventral quadrant of the soma (Figure 1D).

**Figure 1.**
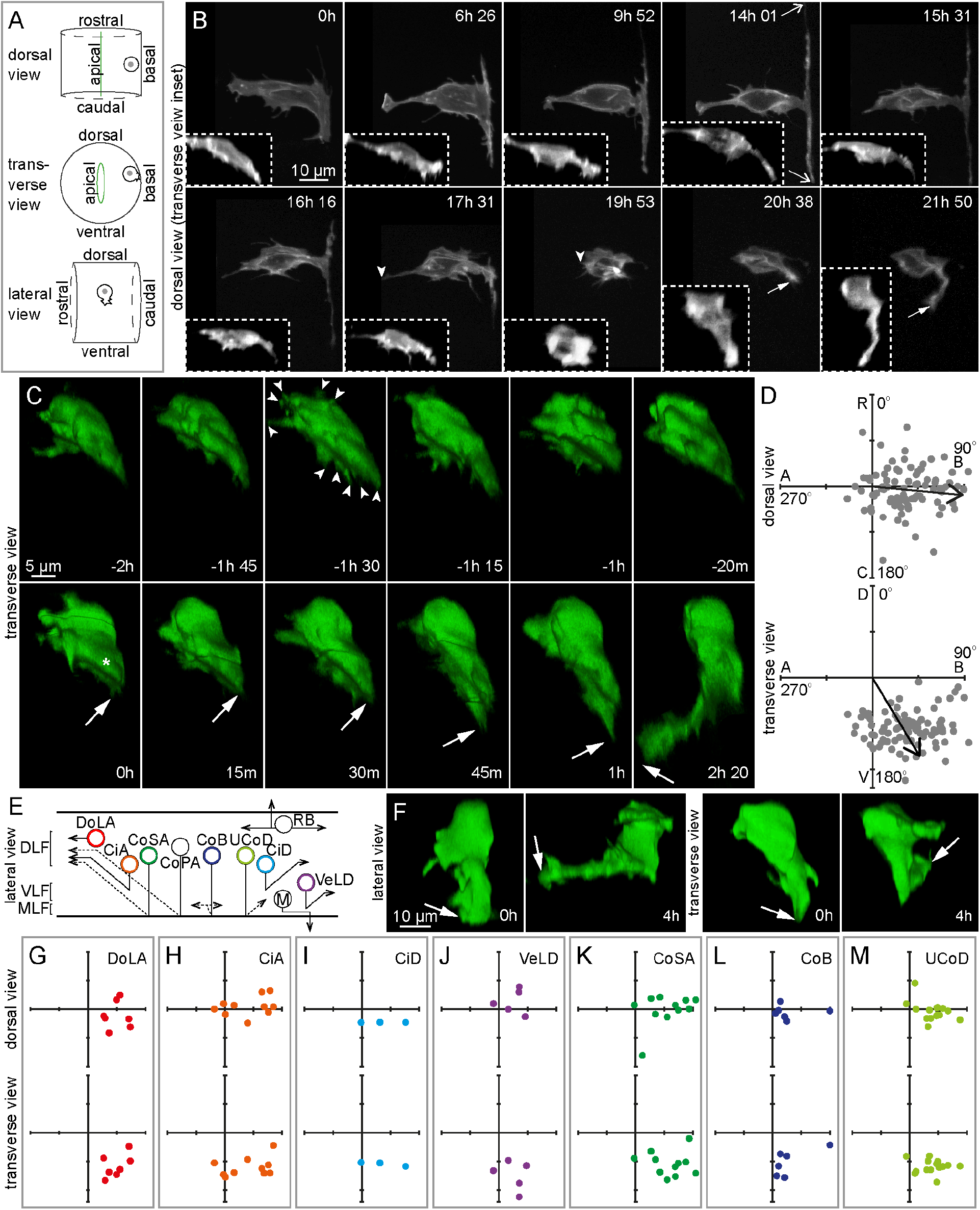
Axon initiation is highly stereotyped and occurs at the baso-ventral aspect of the soma irrespective of subsequent axon trajectory. (**A**) Diagram to illustrate the three different views – dorsal, transverse and lateral – shown in confocal images and 3D reconstructions throughout this paper. (**B**) Image sequence from confocal time lapse shows the early steps in neuronal differentiation. Two transient basal protrusions are extended along the basal surface of the neural tube (6h 26 to 14h 01; open arrows) and then retracted (15h 31 to 17h 31). The apical attachment is also retracted (17h 31 to 19h 53; e.g. −1h 30, arrowheads) before the axon is extended (20h 38 to 21h 50; closed arrows). Images are maximum projections (dorsal views) and transverse reconstructions (transverse views; insets) from confocal z-stacks. (**C**) Image sequence from confocal time lapse shows a neuron before, during and after axon initiation. Prior to axon initiation, the neuron extends multiple small, transient pre-axonal protrusions (−2h to −20m; arrowheads). The nascent axon is extended (0h) and maintained for a short period (0h to 30m) before axon growth begins (45m to 2h 20). Arrows show axon tip. Asterisk marks position of the base of the axon on the soma. Images are transverse reconstructions from confocal z-stacks. (**D**) Plots showing axon position on the soma (e.g. asterisk in Figure 1C 0h) relative to the cell centroid at 0,0 for dorsal and transverse view (n = 86 cells). Axon position is not random (dorsal view *P* < 0.001; transverse view *P* < 0.001). Arrows show mean angle (dorsal view mean = 95.3°; transverse view mean = 148.9°). Data analysed using Moore’s modification of the Rayleigh’s test. R, rostral; C, caudal; A, apical; B, basal; D, dorsal; V, ventral. (**E**) Diagram showing neuronal subtypes in the zebrafish embryos spinal cord. DLF, dorsal longitudinal fasciculus; VLF, ventral longitudinal fasciculus; MLF, medial longitudinal fasciculus; DoLA, dorsolateral ascending; CiA, circumferential ascending; CoSA, commissural secondary ascending; CoPA, commissural primary ascending; CoB, commissural bifurcating; CoD, commissural descending; RB, Rohon-Beard; M, motor; VeLD, ventral longitudinal descending. (**F**) Lateral and transverse reconstructions of DoLA neuron at the time of axon initiation (0h) and during axon growth (4h). Arrows show axon tip. (**G-M**) Plots showing axon position on the soma relative to cell centroid at 0,0 in dorsal and transverse view for DoLA (G; n = 7 cells), CiA (H; n = 10 cells), CiD (I; n = 3 cells), VeLD (J; n = 5 cells), CoSA (K; n = 11 cells), CoB (L; n = 6 cells) and CoD (M; 15 cells) neuronal subtypes.

Several different projection neuron subtypes arise during the first few hours of neurogenesis in the embryonic zebrafish spinal cord (Bernhardt et al., 1990; Hale et al., 2001). They can be distinguished by the dorso-ventral position of the soma within the spinal cord together with their axon trajectory (Figure 1E). Our cell labelling method randomly targeted all of the early embryonic neuronal subtypes reported previously (Figure 1 – figure supplement 2A; (Bernhardt et al., 1990; Hale et al., 2001)). Of the 86 neurons that we analysed for the site of axon initiation (Figure 1D), we were able to classify 53 by neuronal subtype (Figure 1E, F and Figure 1 - figure supplement 2B-G). The axons of many neuronal subtypes grow ventrally and circumferentially before projecting either to the contralateral side of the spinal cord (e.g. CoSA; Figure 1 – figure supplement 1E), or ipsilaterally (e.g. circumferential ascending (CiA) neurons; Figure 1 – figure supplement 1B). We find the site of nascent axon formation was baso-ventral for all of these neuronal subtypes (Figure 1H-M). The only neuronal subtype whose axons are not circumferential are dorsal lateral ascending (DoLA) neurons, which project their axons rostrally towards the hindbrain (Figure 1E; (Bernhardt et al., 1990)). Surprisingly however, DoLA neurons also had a baso-ventral location for their nascent axon (Figure 1F transverse view 0h, G; Video 03). Time lapse imaging showed that, after basolateral nascent axon formation, the growth cone turned and grew rostrally to establish its characteristic axon trajectory (Figure 1F lateral view 4h; Video 03).

Altogether, this confirms and quantifies our previous observation that axon initiation occurs directly from the neuronal soma (Hadjivasiliou et al., 2019). The first persistent axonal protrusion, which we define as the nascent axon, is formed at an extremely stereotyped position such that all analysed neuronal subtypes exhibited a strong baso-ventral bias of the position of nascent axon outgrowth. That this is consistent regardless of the subsequent axonal trajectory shows that axon initiation in the zebrafish spinal cord is a separate process that can be decoupled from axonal growth and guidance.

### Pard3 and Cdh2 are not detected at the growth cone

Several polarity proteins localise to the growth cone in cell culture and have been implicated in axon specification, including Pard3 (Shi 2003) and Cdh2 (previously N-cadherin; (Gärtner et al., 2012)). To investigate the localisation of Pard3 during axon initiation we used a zebrafish Pard3-GFP reporter line (TgBAC(Pard3-GFP)). Time-lapse Airyscan imaging showed clear Pard3-GFP rings at the apical surface of the neural tube, as expected (Figure 1 – figure supplement 3A; (Tawk et al., 2007)). However, we found no evidence for Pard3 localisation in the baso-ventral domain of the soma or in the nascent axon at the time of axon initiation (Figure 1 – figure supplement 3A). We also could not detect any Pard3-GFP signal in the growth cones of growing axons as they crossed the midline (Figure 1 – figure supplement 3C, D). To confirm this result, cells labelled with a fluorescent nuclear marker were transplanted from TgBAC(Pard3-GFP) embryos to wildtype embryos. But even without Pard3GFP expression in surrounding cells, Pard3-GFP could not be detected in the basoventral domain or nascent axons of transplanted cells (Figure 1 – figure supplement 3B).

We performed similar experiments using a zebrafish reporter line for Cdh2 (Tg(cdh2:cdh2-tFT)). We observed clear Cdh2 rings at the apical surface as expected (Symonds et al., 2020) but no accumulation of Cdh2 related to the nascent axon (Figure 1 – figure supplement 3E) or in the growth cones of growing axons as they crossed the midline (Figure 1 – figure supplement 3F).

### Centrosome behaviour prior to axon initiation

Previous studies have suggested that centrosome position is important for positioning the axon (de Anda et al., 2005; Andersen and Halloran, 2012). To get a comprehensive analysis of the centrosome leading up to and during axon initiation we monitored centrosome position from its initial location at the apical surface of the neuroepithelium through to the time of nascent axon establishment. We first focussed on the time when the neuron detaches from the apical surface (see Figure 1B, 17h31 to 19h 53). In *ex ovo* chick neural tube slices the centrosome remains at the apical surface during neuronal differentiation until abscission of the apical processes, when the centrosome is retracted back to the soma along with the apical process. The retracting process abscises from the apical endfoot, which is left behind at the apical surface of the neural tube along with the cilium (Das and Storey, 2014). We too found that the centrosome in zebrafish spinal cord cells is retracted along with the apical process (Figure 2A). However, unlike the chick spinal cord, most zebrafish spinal neurons do not show any apical abscission events (Figure 2B; Video 04). We next used a zebrafish transgenic line with GFP-tagged cilia (Tg(arl13b-GFP)) and observed cilia in the zebrafish spinal cord during a period when we know that many neurons will be retracting their apical processes. We saw many examples of cilia moving from the apical surface to close to the basal surface (Figure 2C), reminiscent of apical retraction. In several instances we could follow a particular cilium continually for up to 45 minutes before it left the apical surface and then moved basally (n = 13 cilia). Cilium length did not change either while at the apical surface or while moving towards the basal surface (Figure 2E). These data show that in zebrafish the cilia are retained by most spinal neurons during apical retraction rather than being abscised and regrown. Finally, we observed centrosomes and cilia at the same time by labelling centrosomes in the zebrafish cilium line. Time lapse videos showed the centrosome and cilia stay in close proximity both at the apical surface and as they moved together towards the basal surface (Figure 2D; Video 05). Immunohistochemistry also showed cilia and centrosomes close together in several positions along the apico-basal axis of the spinal cord, and close proximity was maintained no matter where along they were along this axis (Figure 2F). Altogether this data shows that the centrosome and cilia do not physically dissociate but are retracted together within the apical process during the large majority of neuronal differentiation events in the zebrafish spinal cord. Since it travels in the apical pole of the retracting process, the centrosome locates to the apical pole of the neuronal soma at the end of this phase of differentiation.

**Figure 2.**
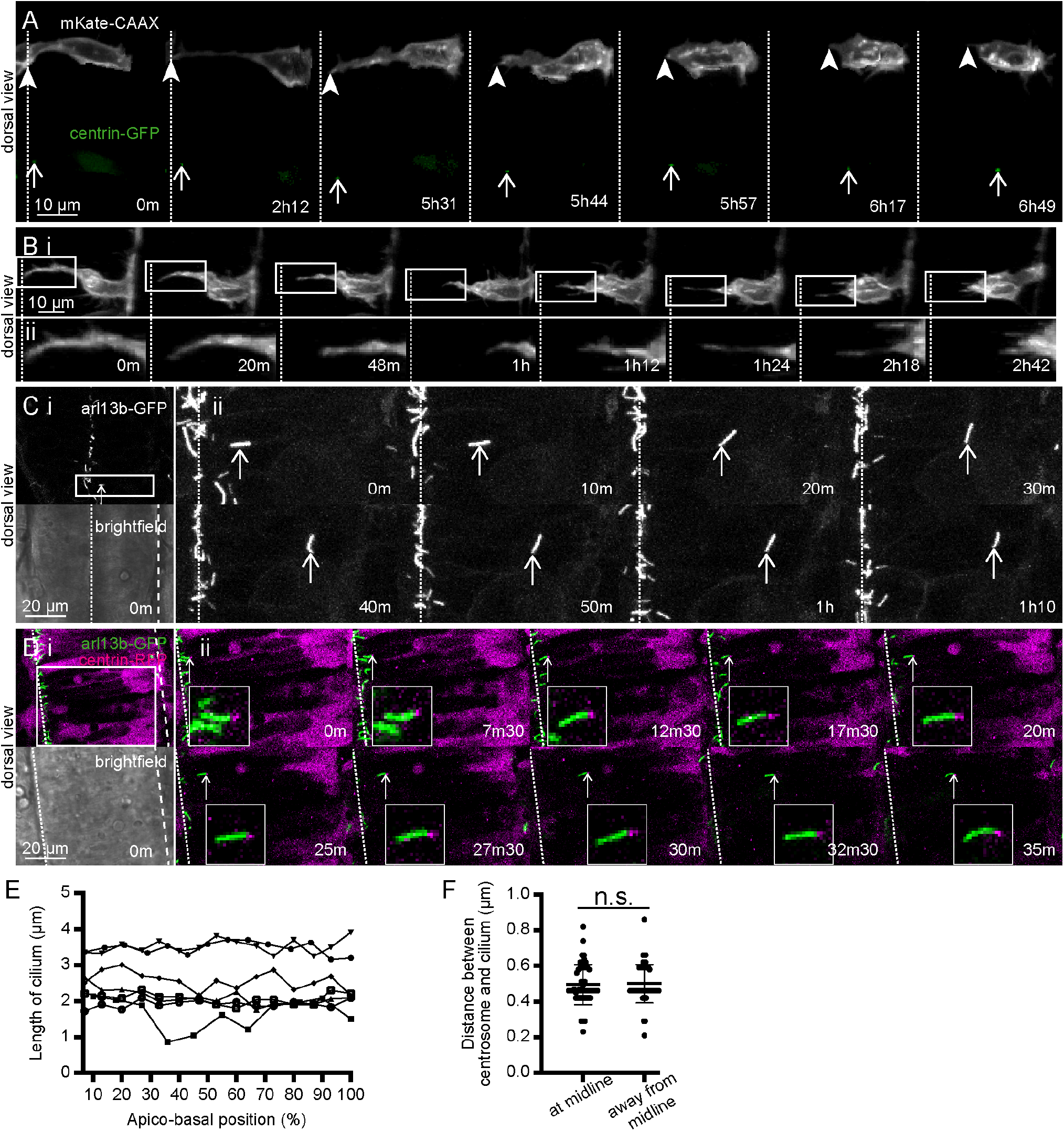
The centrosome and cilium are retracted to the soma during apical process retraction. (**A**) Image sequence showing centrosome position of two adjacent neurons labelled with membrane (grey) and centrosome (green) labels during apical process retraction. The centrosome can be retracted ahead of the apical process (top cell) or stay close to the end of the apical process during retraction (bottom cell). Open arrows = centrosome positions. Arrow heads = tip of apical process. Dotted line = apical surface. Images are maximum projections from confocal z-stacks. (**B**) (i) Image sequence showing a neuron labelled with a membrane marker during apical retraction without any observable abscission event (n = 64/72 cells). (ii) High resolution images of boxed section in (i). Dotted line = apical surface. Images are maximum projections from confocal z-stacks. (**C**) (i) Low magnification overview of the spinal cord of a cilium reporter line, Tg(arl13b-GFP) from a single confocal slice. (ii) High resolution image sequence of boxed section in (i). One GFP-labelled cilium (arrows) moves from apical surface towards the basal surface of the spinal cord (n = 13 cilia). Dotted line = apical surface; dashed line = basal surface. Images are maximum projections from confocal z-stacks. (**D**) (i) Low magnification overview of the spinal cord of a Tg(arl13b-GFP) embryo (green) labelled with centrin-RFP (magenta) from a single confocal slice. (ii) Image sequence of boxed section in (i). A cilium (green) and centrosome (magenta; arrows) move together from apical surface towards the basal surface of the spinal cord (n = 2 cells). Insets show high magnification of boxed section in (ii). Dotted line = apical surface; dashed line = basal surface. Images are maximum projections from confocal z-stacks. (**E**) Graph showing length of cilium as it moves from apical surface (0%) to close to basal surface (100%. Cilium length did not change (*P* > 0.05 for n = 6/7 cilia; non-linear regression). (**F**) Distance between centrosome and cilium in Tg(arl13b-GFP) embryos fixed and processed for immunohistochemistry against GFP to label the cilium and γ-tubulin to label the centrosome. No difference was found in the distance between the two organelles when close to the apical surface or away from the midline (n = 50 cells per condition; *P* = 0.8279; midline mean = 0.4956, s.d = 0.1125; away from midline mean = 0.5004, s.d = 0.1077; Student’s unpaired t-test).

### The centrosome is located on the opposite side of the cell to the nascent axon

As apical retraction was completed the centrosome was close to the apical pole of the neuronal soma. To assess whether proximity of the centrosome is involved in axon initiation we next analysed centrosome position in neurons during establishment of the nascent axon. Time lapse imaging showed that the centrosome was not close to the position of the nascent axon (Figure 3A 0m; Video 06). When the nascent axon is first identifiable, the mean distance between the centrosome and nascent axon was 10.1 μm ± 3.3 (Figure 3C). To put this into perspective, the mean diameter of these cells’ nuclei was 7.8 μm (Figure 3C dotted line, s.d. = 0.7, n = 5 cells). To quantify the spatial relationship between centrosome and nascent axon we analysed their positions with respect to the cell centroid at the time of axon initiation. The centrosome position was highly biased towards the apico-dorsal side of the cell (Figure 3F green dots), placing it on the opposite side of the cell to the baso-ventral site of the nascent axon (Figure 3F grey dots). Further, paired analysis of the positions of the centrosome and nascent axon in the same cell showed that these were different (Figure 3F). Finally, we analysed the slope of vectors linking the centrosome and nascent axon of each cell. The centrosome-axon axis was strongly oriented from apical to basal in the dorsal view and from apico-dorsal to baso-ventral in the transverse view (Figure 3F). These results show clearly that the centrosome is not close to the site of the nascent axon in zebrafish spinal cord neurons *in vivo*; indeed, it is normally on the opposite side of the cell. The centrosome is deposited apically and dorsally following apical retraction and remains in that quadrant of the neuron until after axon initiation.

**Figure 3.**
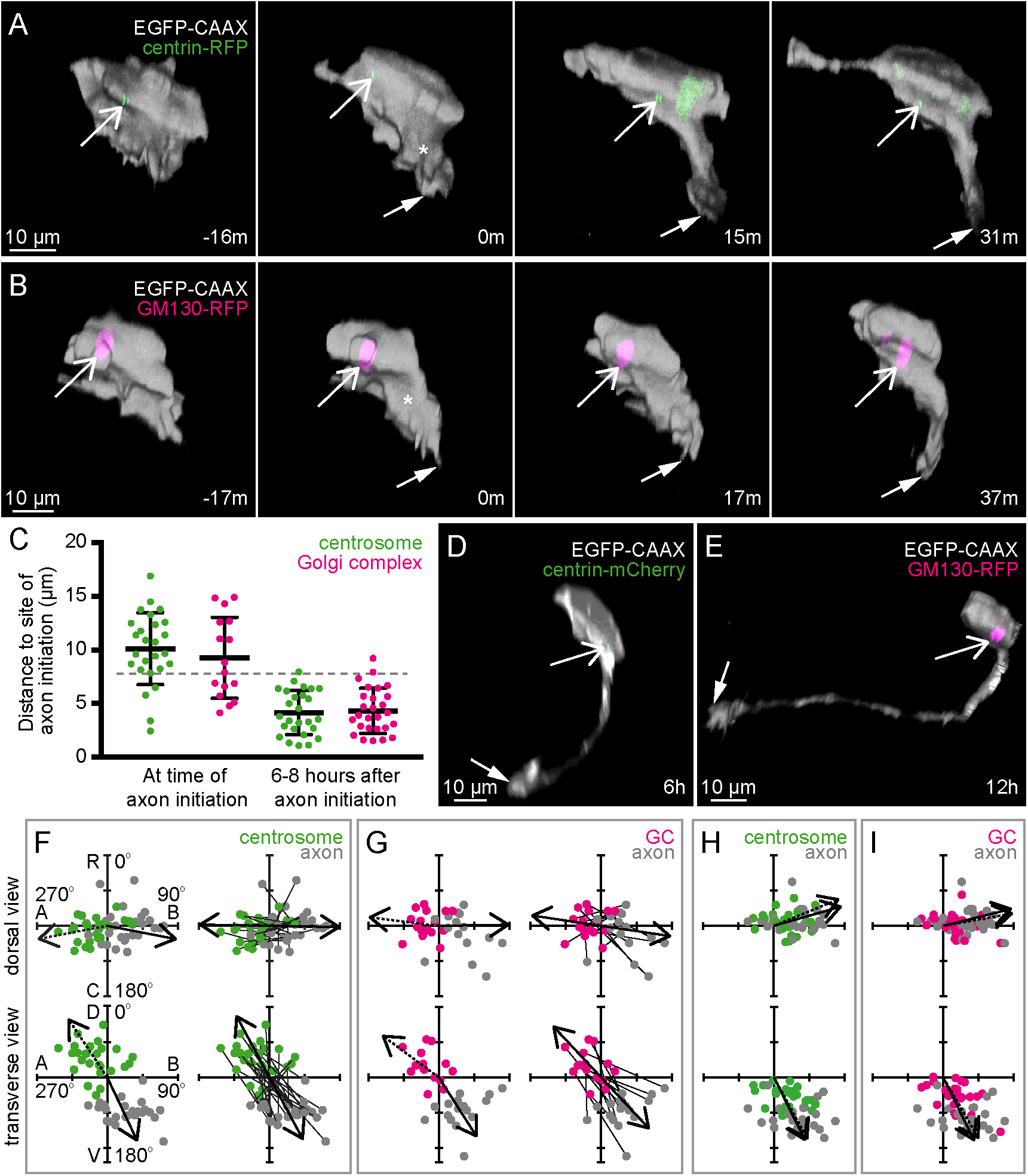
MTOCs are positioned on the opposite side of the cell to the nascent axon. (**A**) Image sequence from confocal time lapse shows a neuron labelled with membrane (grey) and centrosome (green) markers before (−16m), during (0m) and after axon initiation (15m, 31m). The centrosome (open arrows) is located on the opposite side of the cell to the nascent axon (closed arrows). Images are transverse reconstructions from confocal z-stacks. (**B**) Image sequence from confocal time lapse shows a neuron labelled with membrane (grey) and Golgi complex (magenta) markers before (−17m), during (0m) and after axon initiation (17m, 37m). The Golgi complex (open arrows) is located on the opposite side of the cell to the nascent axon (closed arrows). Images are transverse reconstructions from confocal z-stacks. (**C**) Graph showing distance between centrosome (green) or Golgi complex (magenta) and base of axon at time of axon initiation and 6-8 hours after axon initiation. At time of axon initiation: centrosome-axon mean = 10.1 um, s.d. = 3.3, n = 26 cells; Golgi complex-axon mean = 9.3 um, s.d. = 3.8, n = 16 cells. At 6-8 hours after axon initiation: centrosome-axon mean = 4.2 um, s.d. = 2.1, n = 26 cells; Golgi complex-axon mean = 4.3 um, s.d. = 2.1, n = 27 cells. (**D**) Transverse reconstruction from confocal time lapse of a neuron labelled with membrane (grey) and centrosome (green; open arrow) markers during axon growth. Closed arrow = axon tip. (**E**) Transverse reconstruction from confocal time lapse of a neuron labelled with membrane (grey) and Golgi complex (magenta; open arrow) markers during axon growth. Closed arrow = axonal growth cone. (**F**) Plots showing the positions of the centrosome (green) and base of the axon (grey) at the time of axon initiation relative to the cell centroid at 0,0 for dorsal and transverse view (n = 26 cells). Left-hand plots: centrosome position is not random (dorsal view *P* < 0.001; transverse view *P* < 0.001) and dotted arrows show mean angle of centrosome (dorsal view mean = −100.8°; transverse view mean = −33.7°); axon position is not random (dorsal view *P* < 0.001; transverse view *P* < 0.001) and solid arrows show mean angle of axon (dorsal view mean = 101.0°; transverse view mean = 154.8°; Moore’s modification of Rayleigh’s test). Centrosome and axon positions are significantly different (dorsal view 0.001 > *P*; transverse view 0.001 > *P*; Moore’s test for paired data). Right-hand plots: lines connect centrosome and nascent axon from the same cell. Double-headed arrows show average slope of vectors linking centrosome and nascent axon, which are not random (dorsal view *P* < 0.001, mean = 91.0°; transverse view *P* < 0.001, mean = 151.7°; Moore’s modification of Rayleigh‘s test). R, rostral; C, caudal; A, apical; B, basal; D, dorsal; V, ventral. (**G**) Plots showing the positions of the Golgi complex (magenta) and base of the axon (grey) at the time of axon initiation relative to the cell centroid at 0,0 for dorsal and transverse view (n = 16 cells). Left-hand plots: Golgi complex position is not random (dorsal view 0.01 < *P* < 0.05; transverse view *P* < 0.001) and dotted arrows show mean angle of Golgi complex (dorsal view l[mean = − 83.0°; transverse view mean = −53.4°); axon position is not random (dorsal view *P* < 0.001; transverse view *P* < 0.001) and solid arrows show mean angle of axon (dorsal view mean = 89.1°; transverse view mean = 147.4°; Moore’s modification of Rayleigh’s test). Golgi complex and axon positions are significantly different (dorsal view 0.001 > *P*; transverse view 0.001 > *P*; Moore’s test for paired data). Right-hand plots: lines connect Golgi complex and nascent axon from the same cell. Double-headed arrows show average slope of vectors linking Golgi complex and nascent axon (dorsal view *P* < 0.001, mean = 98.5°; transverse view *P* < 0.001, mean = 156.8°; Moore’s modification of Rayleigh’s test). (**H**) Plots showing the positions of the centrosome (green) and base of the axon (grey) 6-12 hours after axon initiation relative to the cell centroid at 0,0 for dorsal and transverse view (n = 26 cells). Centrosome position is not random (dorsal view *P* < 0.001; transverse view *P* < 0.001) and dotted arrows show mean angle of centrosome (dorsal view mean = 67.7°; transverse view mean = 151.6°). Axon position is not random (dorsal view *P* < 0.001; transverse view *P* < 0.001) and solid arrows show mean angle of axon (dorsal view mean = 74.0°; transverse view mean = 153.7°; Moore’s modification of Rayleigh’s test). Centrosome and axon positions are not significantly different (dorsal view 0.5 < *P*; transverse view 0.1 < *P* < 0.5; Moore’s test for paired data). (**I**) Plots showing the positions of the Golgi complex (magenta) and base of the axon (grey) 6-12 hours after axon initiation relative to the cell centroid at 0,0 for dorsal and transverse view (n = 27 cells). Golgi complex position is not random (dorsal view *P* < 0.001; transverse view *P* < 0.001) and dotted arrows show mean angle of Golgi complex (dorsal view mean = 79.9°; transverse view mean = 149.9°). Axon position is not random (dorsal view *P* < 0.001; transverse view *P* < 0.001) and solid arrows show mean angle of axon (dorsal view mean = 76.0°; transverse view mean = 153.9°; Moore’s modification of Rayleigh’s test). Golgi complex and axon positions are different only in transverse view (dorsal view 0.5 < *P*; transverse view 0.005 < *P* < 0.01; Moore’s test for paired data).

These results for the projection neurons of the spinal cord seem to be at odds with a previous report suggesting the centrosome was close to the site of peripheral axon initiation in primary sensory Rohon-Beard neurons in zebrafish embryos (Andersen 2012). Rohon-Beard neurons extend three axons - ascending, descending, and peripheral. We analysed centrosome position in Rohon-Beard neurons in relation to each of these axons (Figure 3 – figure supplement 1A). In 22 of 23 events analysed, the centrosome was more than 10 μm from the site of axon initiation (Figure 3 – figure supplement 1B). When centrosome and axon positions were assessed with respect to the cell centroid, the centrosome was not close to the site of axon initiation but was located in the apical side of the soma when the ascending and descending axons were initiated, as previously reported (Figure 3 – figure supplement 1C; (Andersen and Halloran, 2012)). When the peripheral axon was initiated, the centrosome was closer to the cell centroid than to the axon for every cell analysed. The rostral-caudal position of the centrosome did not appear to correlate with the rostral-caudal position of the site of peripheral axon initiation (Figure 3 – figure supplement 1C dorsal view), further supporting that the centrosome is not close to the site of peripheral axon during initiation. Thus, although the centrosome moves towards the basal side of Rohon-Beard neurons during peripheral axon initiation and growth (Figure 3 – figure supplement 1A 9h; (Andersen and Halloran, 2012)), it is not close to the site of axon initiation when an axon is first extended.

### The Golgi complex is also located on the opposite side of the cell to the nascent axon

The Golgi complex can also nucleate microtubules (Chabin-Brion et al., 2001) and is close to the neurite that becomes the axon *in vitro* (de Anda et al., 2005). As such, it could potentially act as an alternative MTOC independently of the centrosome. To investigate this, spinal cord cells were randomly labelled with a membrane marker and a Golgi complex marker (GM130-RFP or -GFP). Time-lapse analysis showed the Golgi complex was not close to the site of nascent axon formation (Figure 3B 0m, 3C). Like the centrosome, the position of the Golgi complex was biased towards the apico-dorsal side of the cell (Figure 3G magenta dots), on the opposite side of the cell to the nascent axon (Figure 3G grey dots). The Golgi complex-axon axis was strongly oriented from apical to basal in the dorsal view and from apico-dorsal to baso-ventral in the transverse view (Figure 3G). These results show that the Golgi complex is also not close to the site of axon initiation *in vivo*.

Finally, we used immunohistochemistry to investigate γ-tubulin location. γ-tubulin is highly likely to be required for microtubule nucleation in cells and so marks any potential MTOC (Moritz and Agard, 2001). We could only find obvious γ-tubulin accumulation at one concentrated point in each neuron that appeared to correspond with the centrosome (Figure 3 – figure supplement 2). Along with analysis of centrosome and Golgi positioning, this suggests that there is no potential MTOC close to the site of axon initiation.

### Both centrosome and Golgi move to the base of the axon after its initiation

Some previous studies have shown that the centrosome and Golgi complex are located at the base of axons in cell culture (de Anda et al., 2005; Stiess et al., 2010) and the centrosome moves towards the basal side of the cell during peripheral axon extension in Rohon-Beard neurons (see Figure 3 – figure supplement 1; (Andersen and Halloran, 2012)). We hypothesised that the proximity of these MTOCs to the axon may reflect axon growth rather than axon initiation, so we analysed the position of the centrosome and Golgi complex during axon pathfinding, between six and twelve hours after establishment of the nascent axon (Figure 3D, E). We found that both organelles had moved close to the base of the axon (Figure 3C). The position of all of these organelles was at the baso-ventral side of the cell and was not random (Figure 3H, I). Paired analysis of the positions of the centrosome or Golgi complex and base of the axon in the same cell showed that these were not different in most cases (Figure 3H, I). Altogether our results show MTOCs are closely related to the base of the axon during axon growth but not during axon initiation.

### Actin accumulation precedes enrichment of growing microtubule plus-ends during nascent axon formation

Although the location of MTOCs is not close to the site of nascent axon initiation it is still possible that microtubules are involved in specifying the nascent axon. Alternatively actin accumulation may precede microtubules to specify this site, as has been observed for neurite initiation (Zhang 2016). To identify the earliest cytoskeletal component we investigated the localisation of various cytoskeletal markers in individual neurons before, during and after axon initiation.

A constitutively active version of the kinesin 1 motor domain, Kif5c560, is trafficked specifically on axonal microtubules and is an early axonal marker *in vitro* and *in vivo* (Jacobson et al., 2006; Randlett et al., 2011). We found that Kif5c560-YFP localised basally in neuroepithelial cells (Figure 4A). In neurons it was fairly evenly localized throughout the cell body before nascent axon establishment and was then gradually restricted basally and ventrally in the soma before being enriched in the nascent axon and in the growth cone during axonal growth (Figure 4B and Figure 4 – figure supplement 1; Video 07; n = 10/13 cells). This suggests that, despite the centrosome sitting distant to the nascent axon, microtubule-based traffic is an early contributor to the nascent axon organisation.

**Figure 4.**
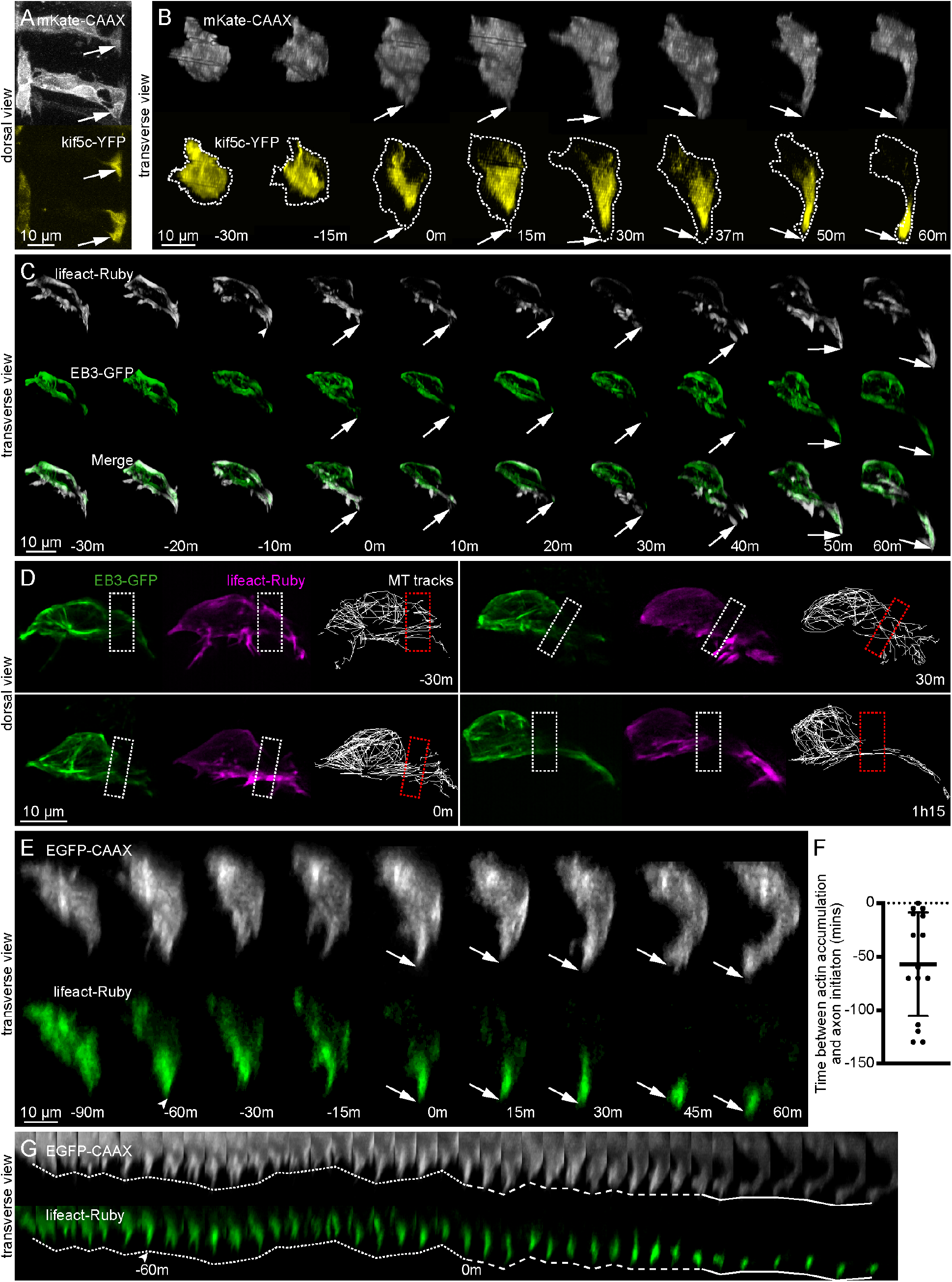
Actin accumulation precedes enrichment of microtubule plus-ends during nascent axon formation. (**A**) Maximum projection from confocal time lapse of neuroepithelial cells expressing a membrane marker (grey) and kif5c560-YFP (yellow). Kif5c560-YFP is enriched in their basal end-feet (arrows). (**B**) Image sequence from confocal time lapse of a neuron labelled with a membrane marker (grey) and Kif5c560-YFP (yellow) before, during (0m) and after axon initiation. Dotted line shows cell outline. Arrows = axon tip. Images are transverse reconstructions from confocal z-stacks. (**C**) Image sequence from confocal time lapse of a neuron labelled with lifeact-Ruby (grey) and EB3-GFP (green) before, during (0m) and after axon initiation. Dotted line shows cell outline. Arrows = axon tip. Images are transverse reconstructions from confocal z-stacks. (**D**) Maximum projections of neuron in (C) showing EB3-GFP (green), lifeact-Ruby (magenta) and microtubule tracks (white) before, during (0m) and after axon initiation. Boxes show transition area between cell body and axonal protrusion. (**E**) Image sequence from confocal time lapse of a neuron labelled with a membrane marker (grey) and lifeact-Ruby (green) before, during (0m) and after axon initiation. Dotted line shows cell outline. Arrowhead = earliest persistent lifeact-Ruby accumulation. Arrows = axon tip. Images are transverse reconstructions from confocal z-stacks. (**F**) Graph showing time (minutes) between actin accumulation and nascent axon initiation (n = 15 cells). (**G**) Kymograph from transverse reconstruction of neuron in (E) showing lifeact-Ruby (magenta) localisation to axon over time. Arrowhead = earliest persistent lifeact-Ruby accumulation. Time of nascent axon initiation is 0m. Dotted line = ventral-most tip of pre-axonal protrusion; dashed line = ventral-most tip of nascent axon; solid line = ventral-most tip of axon during axon growth.

To compare the timing of actin accumulation and the invasion of growing microtubules into the nascent axon we analysed the distribution of filamentous actin (F-actin) and EB3 simultaneously. EB3 is a microtubule plus-end binding protein that labels the tips of all growing microtubules. We mosaically expressed EB3-GFP mRNA to observe growing microtubules along with lifeact-Ruby mRNA to label F-actin-based protrusions before, during and after axon initiation. We found that EB3 was localised throughout the cell before axon initiation (Figure 4C −30m to −10m). In contrast to Kif5c560, however, most EB3 was still located in the soma during nascent axon establishment and only a few microtubule plus-ends were localised within the nascent axon itself (Figure 4C 0h; Video 08). EB3 was subsequently enriched in the axonal growth cone during axon growth (Figure 4C 50m and 60m, and Figure 4 – figure supplement 2 and 3). Close examination of EB3 before, during and after nascent axon formation showed that very few microtubule plus-ends grew from the cell body into the new-born protrusion compared to the amount of growing microtubules in the cell body (Figure 4D; Video 09). Simultaneous imaging of lifeactRuby however showed F-actin is enriched in the nascent axon before EB3-GFP localisation (Figure 4C and Figure 4 – figure supplement 3; n = 6/8 cells). Live imaging of neurons expressing combinations of Kif5c560-YFP, lifeact-Ruby and EB3-GFP during axon initiation supported this observation and showed that lifeactRuby localisation to the nascent axon preceded that of Kif5c560 and EB3-GFP (Figure 4 – figure supplement 1-6, n = 3/4 cells). In fact, co-labelling with a membrane probe and lifeactRuby showed that F-actin was persistently localised baso-ventrally before a persistent nascent axon protrusion (Figure 4E-G and Figure 4 – figure supplement 4; Video 10; n = 14/15 cells). In many cases lifeactRuby was localised baso-ventrally more than 30 minutes before nascent axon protrusion (Figure 4F; n = 10/15 cells). Thus, F-actin accumulates at the site of the nascent axon before it is stabilized and is the earliest cytoskeletal sign of specification of nascent axon formation.

### Nascent axons form in the absence of microtubules

The finding that F-actin accumulation consistently preceded the localisation of an axon-specific microtubule motor such as kif5c560 suggests F-actin accumulation may be the primary cytoskeletal element necessary for axon initiation. To test this we examined axon initiation in the absence of microtubules. We labelled cells for membrane and F-actin and bathed embryos in 5 μg/mL nocodazole for 60-180 minutes to depolymerize microtubules (Head et al., 1985; Jordan and Wilson, 1998; Gallo and Letourneau, 1999). By 30 minutes EB3-GFP labelled comets that label growing microtubules have disappeared from cells (Figure 5 – figure supplement 1B) and 45 minutes after nocodazole addition immunohistochemistry against α- tubulin shows that the whole filamentous microtubule array is completely disrupted in newborn neurons and neuroepithelial cells (Figure 5 – figure supplement 1A). We then analysed whether nascent axons could form from 45 minutes after nocodazole treatment.

We focussed on 54 neurons that had retracted their apical and basal processes and had not yet extended a nascent axon (see Figure 1B, 19h53) prior to nocodazole addition. Most of these cells completely retracted any small protrusions or filopodia upon nocodazole treatment (45/54 cells). However, all cells developed multiple short, thin, transient protrusions during nocodazole treatment (Figure 5A-E); we call these non-axonal protrusions. They were reminiscent of pre-axonal protrusions in WT cells (see Figure 1C −1h 30). Non-axonal protrusions were often extended from multiple locations on the cell, either consecutively or sequentially, but the vast majority protrude from the ventral side of the cell (Figure 5A, G). This suggests cells can still respond to ventral cues in the absence of microtubules. A quarter of the cells developed a *de novo* protrusion with characteristics reminiscent of a nascent axon during nocodazole treatment (Figure 5B and Video 11; 14/54 cells). These protrusions, like nascent axons on normal neurons (see Figure 1D), were predominantly initiated from the baso-ventral quadrant of the cell and were longer and more persistent than non-axonal protrusions, lasting at least 15 minutes or until the end of nocodazole treatment (Figure 5C-F).

**Figure 5.**
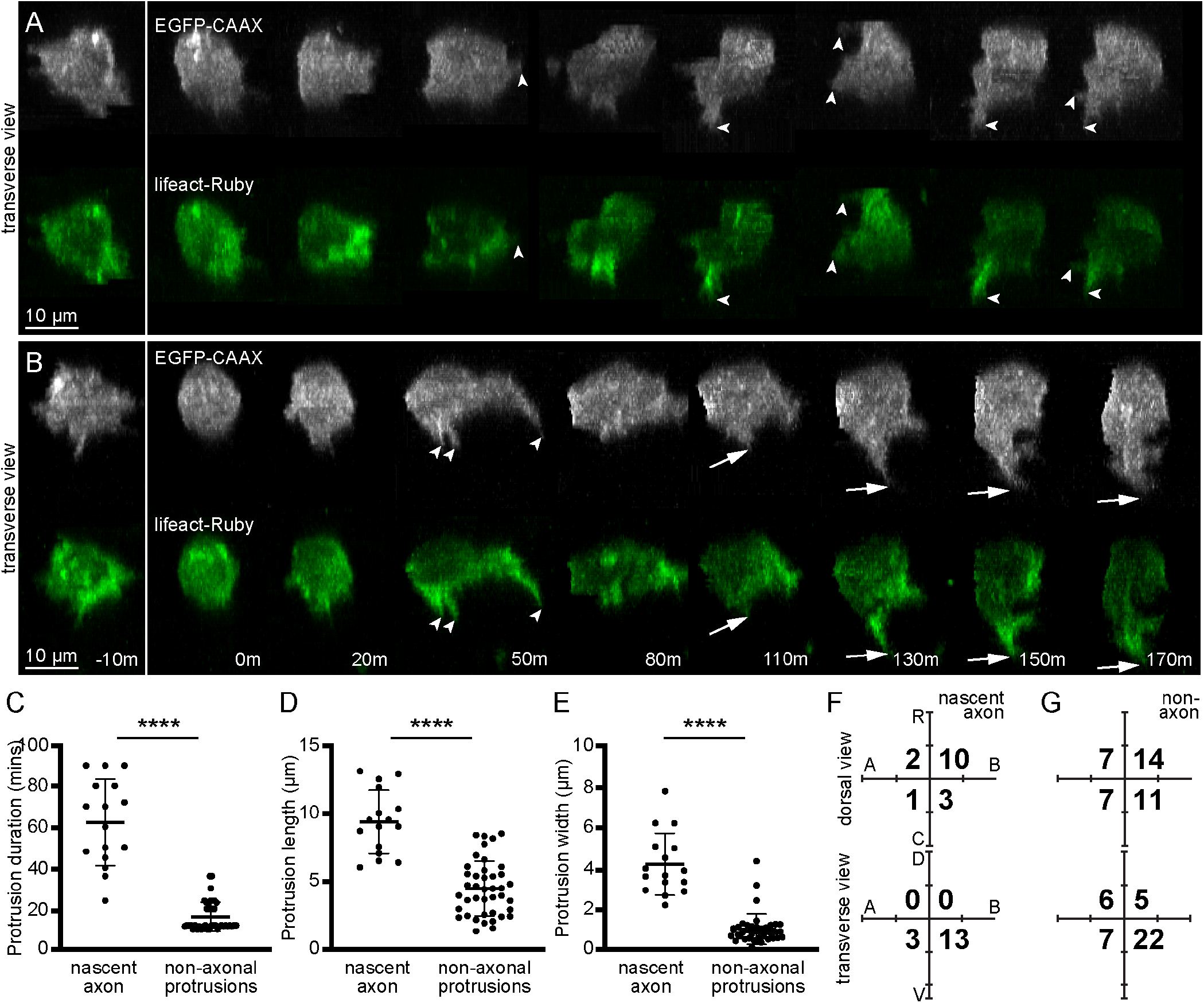
Nascent axons form in the presence of nocodazole, a microtubule polymerisation inhibitor. (**A**) Image sequence from confocal time lapse of a neuron that does not extend a nascent axon during nocodazole treatment labelled with a membrane marker (grey) and lifeact-Ruby (green) before (−10m) and during (0m to 170m) nocodazole treatment. Protrusions present before nocodazole addition are retracted upon nocodazole treatment (20m). Short, transient non-axonal filopodia are extended during nocodazole treatment (arrowheads). Images are transverse reconstructions from confocal z-stacks. (**B**) Image sequence from confocal time lapse of a neuron that extends a nascent axon during nocodazole treatment labelled with a membrane marker (grey) and lifeact-Ruby (green) before (−10m) and during (0h to 170m) nocodazole treatment. Small protrusions present before nocodazole addition (−10m) are retracted upon nocodazole treatment (0m). A nascent axon-like protrusion (long, broad, long-lived) is extended during nocodazole treatment (110m to 170m, arrows). Transient non-axonal protrusions are also present (arrowheads). Images are transverse reconstructions from confocal z-stacks. (**C**) Graph showing duration (minutes) of nascent axon-like protrusions and non-axonal protrusions. Nascent axons: n = 16 protrusions from 16 cells; mean = 62.19 minutes, s.d. = 20.5. Non-axons: n = 41 protrusion from 16 cells; mean = 16.18 minutes, s.d. = 7.15. *P* < 0.0001, Student’s two-tailed t-test. (**D**) Graph showing length (μm) of nascent axon-like protrusions and non-axonal protrusions. Nascent axons: n = 16 protrusions from 16 cells; mean = 9.354 μm, s.d. = 2.356. Non-axons: n = 41 protrusion from 16 cells; mean = 4.407 μm, s.d. = 2.024. *P* < 0.0001, Student’s two-tailed t-test. (**E**) Graph showing width (μm) at the base of nascent axon-like protrusions and non-axonal protrusions. Nascent axons: n = 16 protrusions from 16 cells; mean = 4.196 μm, s.d. = 1.527. Non-axons: n = 41 protrusions from 16 cells; mean = 0.984 μm, s.d. = 0.757. *P* < 0.0001, Student’s two-tailed t-test. (**F**) Plots showing nascent axon position on the soma relative to the cell centroid at 0,0 for dorsal and transverse view (n = 16 cells). Numbers show total count of axons was in each quadrant. R, rostral; C, caudal; A, apical; B, basal; D, dorsal; V, ventral. (**G**) Plots showing the position of non-axonal protrusions on the soma relative to the cell centroid at 0,0 for dorsal and transverse view (n = 41 protrusions from 16 cells). Numbers represent total count of protrusions originating in each quadrant.

These results show microtubules are not required for the initiation of a nascent axon and, together with our results showing that F-actin localisation preceded that of microtubule markers, suggests that F-actin accumulation is the key cytoskeletal element for nascent axon initiation. The retraction of nascent axons that existed before nocodazole treatment suggests microtubules are important stabilising nascent axons in addition to their previously suggested role in axon maintenance and growth (Letourneau and Ressler, 1984; Hahn et al., 2019).

### Laminin provides a basal cue for axon initiation

Finally, we investigated external factors that may be responsible for directing nascent axon formation. As axon initiation occurs from the baso-ventral side of the soma, we hypothesised that the nascent axon is extended adjacent to the neuroepithelial basal surface. We investigated this by randomly labelling cells in a utrophin reporter line that marks the neuroepithelial basal surface (Tg(actb1:*utr-mCherry*); Figure 6A). We used time-lapse imaging with Airyscan acquisition and processing to observe the position of the neuronal soma, the base of the nascent axon and the tip of the nascent axon protrusion with respect to the neuroepithelial basal surface at the time of axon initiation (Figure 6C). Fluorescence intensity analysis showed that all three regions of the neuron were within a few microns of the edge of the neuroepithelium (Figure 6C, E).

**Figure 6.**
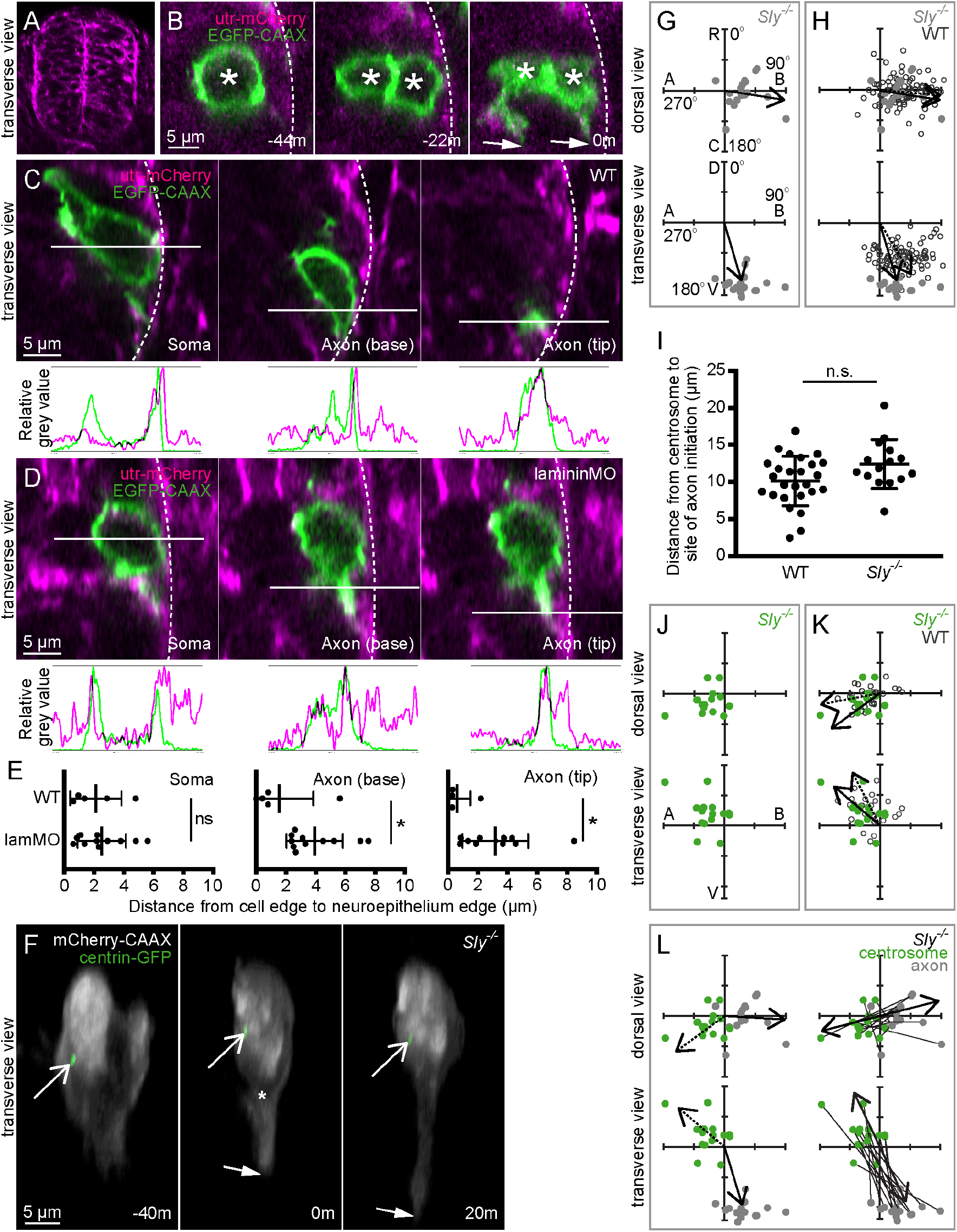
Laminin provides a basal cue for axon initiation. (**A**) Transverse section from a confocal z-stack showing the whole neural tube of a utr-mCherry embryo, showing localisation to the basal surface. (**B**) Transverse sections from a confocal z-stack of a neuron at the time of nascent axon initiation labelled with a membrane marker (green) in a utr-mCherry embryo (magenta) to identify the basal surface of the spinal cord. Three different sections show the middle of the soma, the axon initiation site, and the axon tip. Solid lines show where relative grey values were measured. Dotted lines = basal surface. Green and magenta peaks in graphs show relative positions of cell membrane and basal surface, respectively. (**C**) Image sequence from confocal time-lapse shows one non-apical progenitor (−44m) that undergoes mitosis to produce two neurons (−22m), of which one is not in contact with the basal surface. Both neurons extend nascent axons (0m). Dotted line = basal surface. Asterisks = cell bodies. Arrows = axon tips. Images are transverse sections from confocal z-stacks. (**D**) Transverse sections from confocal z-stacks of a neuron at the time of nascent axon initiation labelled with a membrane marker (green) in a utr-mCherry embryo (magenta) injected with lamMO. Three different sections show the middle of the soma, the axon initiation site, and the axon tip. Solid lines show where relative grey values were measured. Dotted lines = basal surface. Green peaks in relative grey value graphs show edge of cell membrane, while magenta peaks show basal surface. (**E**) Graphs showing distance (μm) between the basal edge of the soma, the axon initiation site or the axon tip and the basal surface of the spinal cord. Measurements were performed by measuring between basal-most green and magenta peaks in graphs of relative grey values (A,C). WT n = 5 cells; lamMO = 11 cells. Soma: WT mean = 2.007 μm, s.d. = 1.055; lamMO mean = 1.871, s.d. = 1.830. Student’s two-tailed test, *P* = 0.880. Axon initiation site: WT mean = 1.366 μm, s.d. = 3.13; lamMO mean = 2.353, s.d. = 2.125. Student’s two-tailed test, *P* = 0.468. Axon tip: WT mean = 0.637 μm, s.d. = 1.078; lamMO mean = 2.903, s.d. = 2.0433. Student’s two-tailed test, *P* = 0.037. (**F**) Image sequence from confocal time lapse shows a neuron in a *Sly*^−/−^ embryo labelled with membrane (grey) and centrosome (green) markers before (−40m), during (0) and after axon initiation (20m). Open arrows = centrosome; closed arrows = axon tip; asterisk = axon position at time of nascent axon initiation. Images are transverse reconstructions from confocal z-stacks. (**G**) Plots showing axon position on the soma relative to the cell centroid at 0,0 for dorsal and transverse view in *Sly*^−/−^ embryos (n = 18 cells). Axon position is not random (dorsal view *P* < 0.001; transverse view *P* < 0.001). Arrows show mean angle (dorsal view mean = 98.8°; transverse view mean = 161.1°; Moore’s modification of Rayleigh’s test). (**H**) Plots showing merge of WT (open circles) and *Sly*^−/−^ (grey dots) axon positions on the cell body relative to cell centroid at 0,0 for dorsal and transverse views. Solid arrows show mean angle from *Sly*^−/−^ embryos (n = 18 cells); dotted arrows show mean angles of WT embryos (n = 86 cells). Axon positions in WT and *Sly*^−/−^ are not significantly different in dorsal view (0.2 < *P* < 0.5) but are in transverse view (P < 0.001; Batschelet’s alternative to Hotelling test). (**I**) Graph showing distance between centrosome and base of axon at time of axon initiation in WT and *Sly*^−/−^ embryos. WT: n = 26 cells, mean = 10.13 μm, s.d. = 3.35. *Sly*^−/−^: n = 15 cells, mean = 12.41 μm, s.d. = 3.281. One-way ANOVA, *P* = 0.123. (**J**) Plots showing merge of WT (open circles) and *Sly*^−/−^ (green dots) centrosome positions on the cell body relative to cell centroid at 0,0 for dorsal and transverse views. Solid arrows show mean angle from *Sly*^−/−^ embryos (n = 15 cells); dotted arrows show mean angles of WT embryos (n = 26 cells). Centrosome positions are not significantly different between WT and <u>*Sly*</u>^−/−^ (dorsal view 0.2 < *P* < 0.2, transverse view 0.1 < *P* < 0.2; Batschelet’s alternative to Hotelling test). (**K**) Plots showing the positions of the centrosome (green) and base of the axon (grey) in *Sly*^−/−^ embryos at the time of axon initiation relative to the cell centroid at 0,0 for dorsal and transverse view (n = 15 cells). Left-hand plots: centrosome position is not random (dorsal view *P* < 0.001; transverse view *P* < 0.001) and dotted arrows show mean angle of centrosome (dorsal view mean = −129.0°; transverse view mean = −57.0); axon position is not random (dorsal view *P* < 0.001; transverse view *P* < 0.001) and solid arrows show mean angle of axon (dorsal view mean = 95.1°; transverse view mean = 168.8°; Moore’s modification of Rayleigh’s test). Centrosome and axon positions are significantly different (dorsal view 0.001 > *P*; transverse view 0.001 > *P*; Moore’s test for paired data). Right-hand plots: lines connect centrosome and nascent axon from the same cell. Double-headed arrows show average slope of vectors linking centrosome and nascent axon, which are not random (dorsal view *P* < 0.001, mean = 66.1°; transverse view *P* < 0.001, mean = 150.9°).

One neuronal subtype in the zebrafish spinal cord derives from division of a non-apical progenitor (Vsx1 progenitors) close to the basal surface of the neural tube. The Vsx1 progenitors are rare and undergo a final mitosis producing two neurons that rapidly extend axons (McIntosh et al., 2017). We observed two cases of non-apical progenitor divisions in which mitotic cleavage resulted in one daughter adjacent to the basal surface and the other apparently not in contact with the basal surface (Figure 6B). Interestingly, the daughter without contact with the basal surface initiated its axon from the ventral but not basal side of the soma while the daughter adjacent to the basal surface initiated its axon baso-ventrally along the basal surface as expected (Figure 6B 0m; n = 2/2 divisions).

This led us to hypothesise that the basal surface of the neural tube may provide a directional cue for nascent axon specification in spinal cord neurons. The extracellular matrix protein laminin, a component of the basal lamina, can influence neuronal polarity and promote axon outgrowth *in vitro* (Esch et al., 1999), and orient axon outgrowth in zebrafish retinal ganglion cells *in vivo* (Randlett et al., 2011), so it is a good candidate to provide a directional cue. We used utrophin reporter line embryos that had been injected with LamininC1 morpholino (lamMO; (Parsons et al., 2002)) to ask whether the presence of laminin influenced the site of axon initiation. Fluorescence intensity analysis showed that in the Laminin-depleted embryos the site of axon initiation and the tip of the nascent axon protrusion were significantly further away from the edge of the basal lamina than in control embryos (Figure 6D, E). However the position of the soma was not affected, indicating that this is not due to incorrect positioning of the neuron (Figure 6E).

To examine the role of laminin in neuronal polarity more closely we analysed the site of axon initiation in the *Sly/LamC1* zebrafish mutant line, which have no detectable laminin expression in the basal lamina at this embryonic stage (Figure 6F, compare to Figure 1C; Video 12). There was no difference in the position of nascent axon initiation between controls and *Sly*^−/−^ embryos when analysed from the dorsal perspective but, analysis from the transverse perspective showed the nascent was less basal and more ventral compared to controls (Figure 6F 0m, G, H). LamMO-injected embryos had a similar but more severe phenotype than *Sly*^−/−^ embryos. The basal bias of the site of the nascent axon was lost in cells in lamMO-injected embryos and the ventral bias was increased compared to controls (Figure 6 – figure supplement 1A 0m, B, C). These results suggest that Laminin promotes the basal positioning of the nascent axon formation site in zebrafish spinal cord neurons.

We next assessed the position of the centrosome at the time of axon initiation in laminin-deficient embryos. As the position of the centrosome tends to be on the opposite side of the cell to the nascent axon position in wildtype cells (see Figure 3), we hypothesised that the change in the position of the nascent axon observed in laminin-deficient embryos may be mirrored by a change in centrosome position. However, there was no difference in the position of the centrosome in *Sly*^−/−^ embryos (Figure 6J, K; Video 12), in the distance between the centrosome and the site of nascent axon in *Sly*^−/−^ or lamMO-injected embryos compared to controls (Figure 6I and Figure 6 – figure supplement 1D), or in the centrosome-axon axes between *Sly*^−/−^ or lamMO-injected embryos and controls (Batschelet’s alternative to Hotelling test; *Sly*^−/−^ dorsal view 0.1 < *P* < 0.2, transverse view 0.1 < *P* < 0.2; lamMO dorsal view 0.05 < *P* < 0.2, transverse view 0.5 < *P*). Thus, although the centrosome tends to be opposite the nascent axon site in wildtype, changes in axon position resulting from Laminin depletion do not appear to significantly alter centrosome position.

### Hedgehog signalling does not provide a ventral cue for axon initiation

Our results show that laminin biases the nascent axon formation site towards the basal side of the neuronal soma, but the ventral bias was maintained in the absence of laminin. As such, we attempted to identify a dorso-ventral cue that promoted the ventral positioning of the nascent axon. Zebrafish embryos were exposed to inhibitors of various signalling pathways or proteins that are known to have a dorso-ventral gradient in the spinal cord, that are known to be axon guidance cues, or that have been shown to influence neuronal polarity in other systems: BMPs (Lee et al., 1998; Augsburger et al., 1999), FGF (Walicke et al., 1986), Hedgehog (Charron et al., 2003), and septins (Boubakar et al., 2017) (not shown). Of these, only inhibition of Hedgehog signalling via cyclopamine (100 μM; (Taipale et al., 2000; Chen et al., 2002)) showed potentially promising results (not shown). To test this we analysed axon initiation in *Smu*^*b641*−/−^ embryos, which have a mutation in the Smoothened protein that is required for both canonical and non-canonical Hedgehog signalling (Figure 6 – figure supplement 2A; (Barresi et al., 2010)). Analysis of *Smu*^*b641*−/−^ embryos showed that the position of the nascent axon was slightly caudal compared to controls but it retained baso-ventral positioning (Figure 6 – figure supplement 2B, C). Thus, although Hedgehog signalling is important for spinal cord axonal guidance (Charron et al., 2003), it is not required to position nascent axon formation or for the accurate establishment of neuronal polarity. Finally, we assessed the position of the centrosome in *Smu*^*b641*−/−^ embryos. There was no difference in the distance between the centrosome and the site of axon initiation or in the position of the centrosome in *Smu*^*b641*−/−^ embryos compared to controls (Figure 6 – figure supplement 2D-F).

## DISCUSSION

We have investigated the earliest steps in axon formation in spinal projection neurons *in vivo*. Our key findings are:

- an accumulation of F-actin is an early molecular indicator of axon initiation, precedes the generation of a stable nascent axonal protrusion and precedes microtubule accumulation
- nascent axons are still formed in the presence of the microtubule disruptor nocodozole
- axon initiation is extremely stereotyped across different spinal neuron subtypes irrespective of subsequent axon guidance
- MTOCs, the centrosome and Golgi apparatus, are located on the opposite side of the cell to the site of axon initiation but move to the base of the axon during axon growth
- Laminin is not required for axon initiation or early growth but is a positional cue for axon initiation.

The polarisation of individual neurons into axonal and dendritic compartments is critical for correct nervous system development. The mechanisms that may define axon initiation have been studied for several decades but many of the previous works are compromised by studying this process in neurons growing *in vitro* that lack the complex 3-dimensional architecture and molecular environment of *in vivo* settings. Additionally, some *in vitro* studies may describe the repolarisation of neurons that had previously polarised *in vivo* rather than neurons polarising for the first time (reviewed in (Barnes and Polleux, 2009)). Further, axon initiation *in vitro* is the differentiation of a pre-existing neurite to become an axon (Dotti et al., 1988), making it difficult to differentiate between axon initiation and axonal growth (Jiang and Rao, 2005; Barnes and Polleux, 2009). For some existing models of neuronal differentiation *in vivo* it can be difficult to disentangle axon initiation from neurite growth and neuronal migration (Barnes and Polleux, 2009). Our study of spinal neurons *in vivo* overcomes these reservations. The cell bodies of neurons in the zebrafish spinal cord move to the basal surface of the neuroepithelium before delaminating from the apical surface and remain established at the basal surface for several hours before axon initiation (Hadjivasiliou et al., 2019), thus the remodelling of cell polarity in this system is not complicated by polarity changes related to cell migration. Axon formation occurs directly from the cell body, so can be clearly identified and separated from axon growth, and we find it occurs from a stereotyped position in all of the spinal neuron subtypes that we investigated, independently of the subsequent direction of axon projection (except for Rohon-Beard neurons, which elaborate three axons). As such, the zebrafish embryonic spinal cord provides a complex *in vivo* system where we can definitively separate axon initiation from both axonal growth and neuronal migration. We define the first phase of axon initiation as the nascent axon – this is a protrusion that is beginning to take on the characteristics of an axon and would normally become an axon when stabilised by microtubules.

Our principle finding is that the earliest indication of axon initiation is a biased accumulation of F-actin in the baso-ventral quadrant of the neuron cell body. This first coincides with unstable protrusions from the baso-ventral soma and then with a stable protrusion that we term the nascent axon (Figure 4). Although microtubules rapidly invade the nascent axon, we find nascent axons can still be formed in the absence of microtubules (Figure 5). Thus, nascent axon formation from spinal neurons *in vivo* is an F-actin-based protrusion that forms directly from a specific location on the cell body. The generation of a single stereotyped axonal protrusion from spinal neurons *in vivo* is very different from the axon selection process from multiple random neurites seen in the *in vitro* neuronal polarisation models (e.g. (Dotti et al., 1988), and reviewed in (Barnes and Polleux, 2009)) and this very likely reflects the difference in complexity of environmental cues in these two cases. The very stereotyped location of nascent axon formation from the baso-ventral quadrant of spinal neurons suggests this process is strongly influenced by local environmental cues in the early neural tube and we uncover that one of those cues is Laminin (Figure 6), an extracellular protein abundant at the basal surface of the neural tube. Laminin may play a common role in the differentiation early born neurons as it has previously been shown to influence retinal ganglion cell axon initiation in the retina (Randlett et al., 2011) and axonal growth in the primary sensory Rohon-Beard neurons in zebrafish neural tube (Andersen and Halloran, 2012).

Although nascent axons can be produced in the absence of microtubules, they are not stable and retract (Figure 5B) demonstrating a requirement for microtubules to stabilize this protrusion. Perhaps in this respect microtubules cooperate with dynamic actin in similar ways as they do to facilitate turning in neuronal growth cones (reviewed in (Geraldo and Gordon-Weeks, 2009)), and to promote axon specification (Bradke and Dotti, 1999; Geraldo et al., 2008; Witte et al., 2008; Zhao et al., 2017) and neurite initiation *in vitro* (Dent et al., 2007; Flynn et al., 2012). Increased numbers of microtubules and enriched microtubule plus-ends could play an important role in anterograde transport (reviewed in (Schelski and Bradke, 2017)). However, in spinal neurons *in vivo* we see few EB3-labelled growing microtubule plus-ends in both pre-axonal protrusions and in the nascent axon, illustrating that there are few growing microtubules until axonal growth commences.

Consistent with our observation that microtubules are not required for nascent axon formation, we also show neither the centrosome nor the Golgi complex is close to the site of axon initiation in spinal neurons. Although several studies suggest the centrosome and Golgi complex are close to the base of the neurite that becomes the axon (Zmuda and Rivas, 1998; de Anda et al., 2005), our results support previous observations that show centrosome proximity is not required for axon initiation (Dotti and Banker, 1991; Zolessi et al., 2006; Distel et al., 2010; Gärtner et al., 2012). There are several potential explanations for discrepancies between these findings. There may be innate differences in cytoskeletal organisation between different neuronal subtypes or species; differences in substrate properties may alter cytoskeletal organisation, as has been shown for migrating cells (Pouthas et al., 2008); or, alternatively, studies showing MTOCs close to the base of the axon may be looking after axon initiation. This is supported by our observation that both the centrosome and Golgi complex move close to the base of the axon during pathfinding and that the centrosome is not close to the site of axon initiation in zebrafish retinal ganglion cells, in which axon initiation can also be easily identified (Zolessi et al., 2006). The centrosome has previously been described as being associated with peripheral axon formation in Rohon-Beard neurons and to be important for its growth (Andersen and Halloran, 2012), but we find that the centrosome is not close to the base of the axon at the time of initiation of any Rohon-Beard axon, including the peripheral axon. Combining these findings and others (e.g. (Stiess et al., 2010)) suggests that centrosome proximity is not required for axon initiation but is required for axon growth, at least at early stages.

Although not close to the site of axon initiation the centrosome is not positioned randomly in spinal neurons; instead, it is consistently opposite the site of axon initiation. As the centrosome is situated apically while the new neuron is still attached to the apical surface and is retracted into the neuronal cell body upon delamination, it may be that its medial position in the soma at axon initiation is simply related to the location of the retracting apical process. Interestingly we found no evidence that apical abscission is required for apical process retraction in the zebrafish spinal cord. This is in contrast to chick and mouse but in agreement with some observations in the zebrafish retina (Zolessi et al., 2006; Das and Storey, 2014; Lepanto et al., 2016). Although the centrosome is not close to the site of nascent axon initiation, it could still play a role in its stabilisation and transition to a growing axon. A small number of microtubules do enter the nascent axon and it is possible that only a few stable microtubules are required for the next steps in axon differentiation. Interestingly, modelling has shown that stochastic microtubule dynamics can lead to stabilisation of the longest microtubules (Seetapun and Odde, 2010), suggesting a method by which the distant centrosome may stabilise the nascent axon on the opposite site of the cell where only the longest microtubules can reach.

The position of the nascent axon is influenced by the extracellular matrix protein laminin at the basal surface of the neural tube, as the loss of laminin leads to the loss of the basal bias to the nascent axon position. In the retina, laminin stabilises newly initiated axons and promotes axonal growth (Randlett et al., 2011). The neurons that we observed were among the earliest that differentiated, meaning that they were almost always already adjacent to the laminin-rich basal surface when extending an axon. It would be interesting to compare this with later-born neurons, which would have earlier-born neurons between them and the basal surface. Nonetheless, we show that axon initiation, stabilisation and growth can occur robustly in the spinal cord without laminin. Although we were unable to find evidence of a dorso-ventrally oriented cue that was required for ventral directional bias of the nascent axon, we can’t rule out that one exists. An alternative to a molecular cue that directs the ventral bias of the nascent axon could be the overall physical architecture of the cells in the early neural tube. All neuroepithelial cells and neurons have a curved morphology in transverse sections, with the lateral poles of both neuroepithelial progenitors and neurons curving ventrally as they approach the basal surface (see Figure 1B, transverse inserts). It seems possible this morphological organisation of cells could provide a 3-dimensional physical substrate or orientation that encourages ventral growth of nascent axon protrusions.

## Acknowledgements

R.E.M. and J.C. were supported by the Wellcome Trust (102895/Z/13/Z). S.P. was supported by an MRC studentship. Thanks to Christopher Rookyard for the image in Figure 6A, Simon Hughes’ lab for kindly providing the *Smu*^*b641*^ zebrafish line, Corinne Houart’s lab for kindly providing the α-tubulin antibody, and to past and present members of the Clarke lab and CDN for constructive feedback and discussion.

R.E.M. and J.C. devised experiments and wrote the manuscript. R.E.M, S.P. and C.A. performed experiments and analysed results.

