## Supplemental Figures for "Actin-based protrusions lead microtubules during stereotyped axon initiation in spinal neurons *in vivo*"

Figure 1 - figure supplement 1

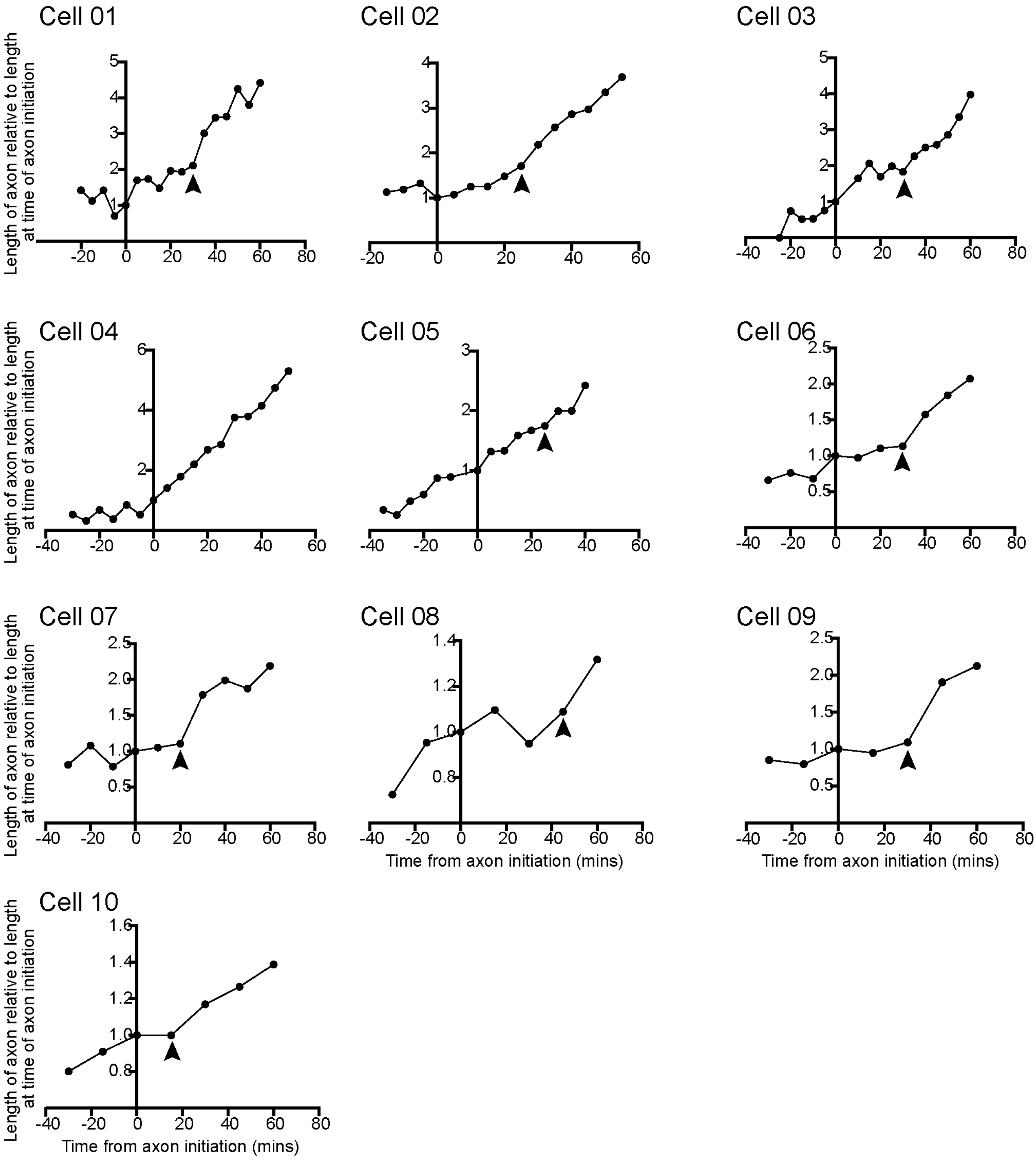

**Figure 1 – Figure Supplement 1.**
Graphs showing the maximum protrusion length from seven different cells before, during (0 mins) and after axon initiation. Length is shown relative to length at the time of axon initiation (0 mins). Arrowheads show transition from nascent axon to growth phase (9/10 cells).

Figure 1 - figure supplement 2

A

| | |
|---|---|
| RB | 29 |
| DoLA | 9 |
| CoPA | 5 |
| CoSA | 28 |
| CiA | 19 |
| CiD | 19 |
| CoB | 14 |
| VeLD | 4 |
| M | 1 |
| UCoD | 24 |

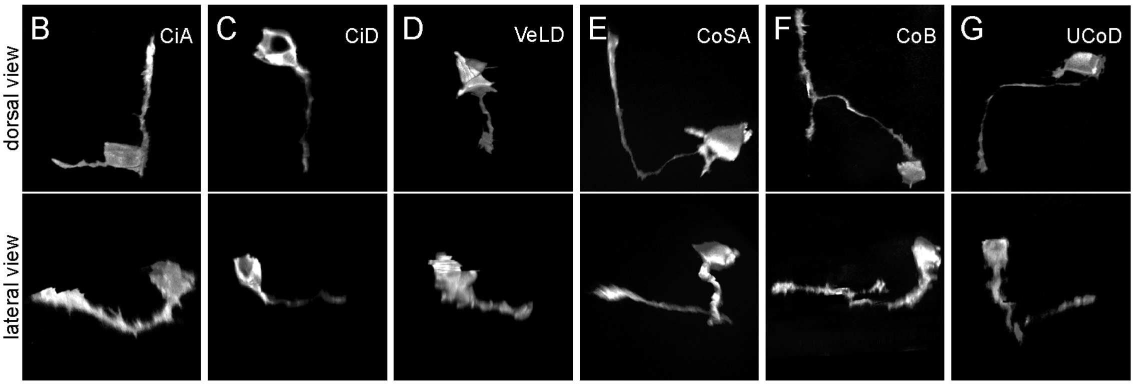

**Figure 1 – Figure Supplement 2.**
(**A**) Numbers of each neuronal subtype labelled. (**B-G**) Examples of CiA (B), CiD (C), VeLD (D), CoSA (E), CoB (F) and CoD (G) neurons and their axon trajectories as dorsal and lateral reconstructions from confocal z-stacks.

Figure 1 - figure supplement 3

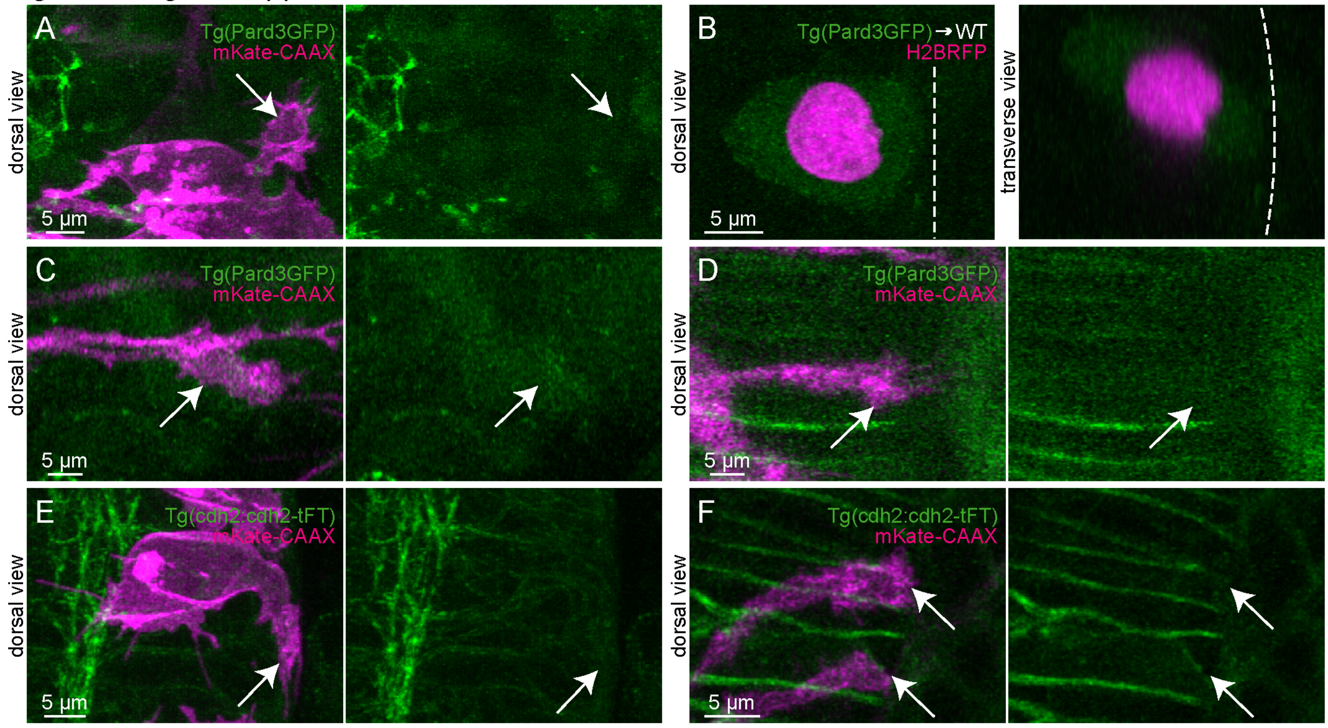

**Figure 1 – Figure Supplement 3.**
(**A**) Maximum projection from one confocal z-stack of a Pard3-GFP transgenic embryo (green). The same field is shown with and without cells labelled with membrane marker (magenta). A nascent axon (arrows) shows no Pard3-GFP accumulation (n = 6/6 cells). Pard3-GFP localises to apical endfeet of neuroepithelial cells on the left of the image. (**B**) Maximum projection of a confocal z-stack (dorsal view) and maximum projection of a reslice of a confocal z-stack (transverse view). A neuron from a Pard3-GFP embryo (green) with a nuclear label (magenta) transplanted to a wildtype embryo shows no Pard3-GFP accumulation (n = 3/3 cells). Dotted lines = basal surface. (**C, D**) Maximum projections of two different confocal z-stacks from Pard3-GFP transgenic embryos (green). Each is shown with and without membrane-labelled axons (magenta) whose growth cones (arrows) are crossing the ventral midline and show no Pard3-GFP accumulation (n = 5/5 cells). Pard3-GFP is expressed in notochord cells ventral to the neural tube in (D). (**E**) Maximum projection from one confocal z-stack of a cdh2-GFP transgenic embryo (green), with and without membrane-labelled cells (magenta). A nascent axon (arrow) shows no enrichment of cdh2-GFP (n = 17 cells). Cdh2-GFP localises to apical endfeet of neuroepithelial cells on the left of the image. (**F**) Maximum projection of a confocal z-stack from cdh2-GFP transgenic embryo (green) with and without membrane-labelled axons (magenta). Growth cones (arrows) are crossing the ventral midline and show no cdh2-GFP enrichment (n = 11 cells). Cdh2-GFP is expressed in notochord cells ventral to the neural tube.

Figure 3 - figure supplement 1

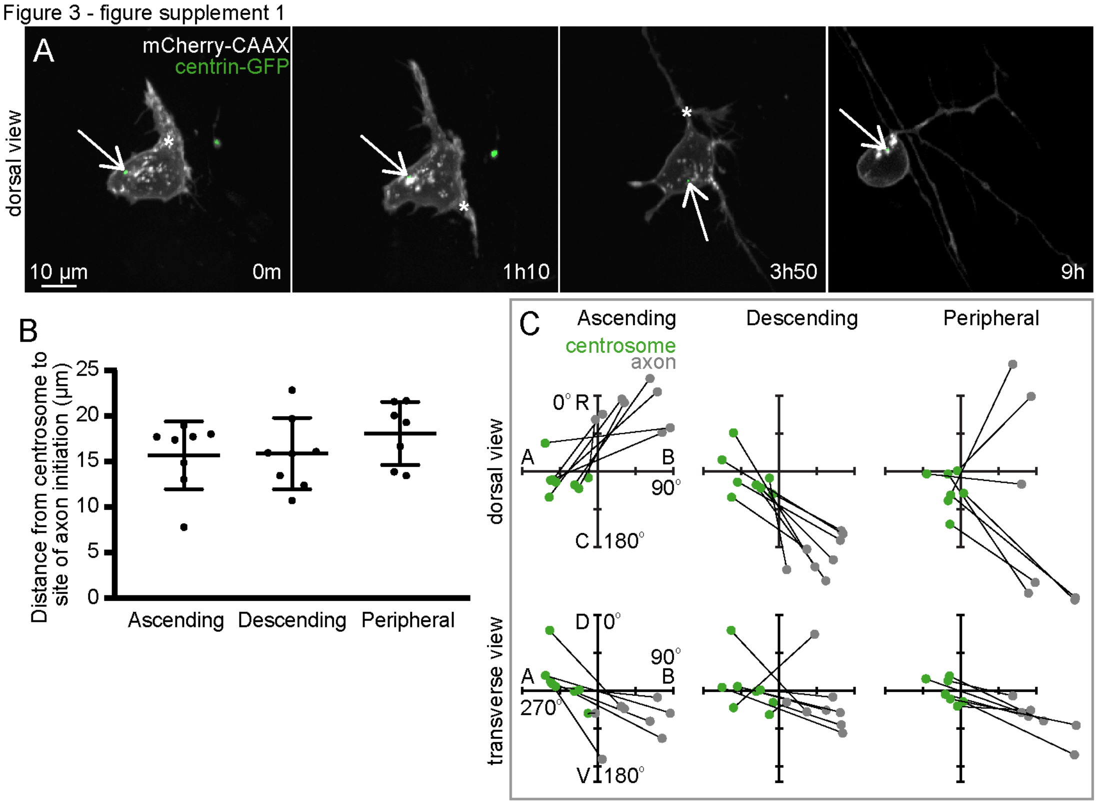

**Figure 3 – Figure supplement 1.**
(**A**) Image sequence from confocal time lapse shows a Rohon-Beard neuron labelled with membrane (grey) and centrosome (green) markers during the initiation of the ascending (0m), descending (1h10) and peripheral axons (3h50), and during axon pathfinding. The centrosome (arrows) is located away from the base of each axon (asterisks) but moves close to the peripheral axon during pathfinding. Images are maximum projections from confocal z-stacks. (**B**) Graph showing distance between centrosome and base of the axon at time of initiation of each axon (n = 23 events from 8 cells). (**C**) Plots showing the positions of the centrosome (green) and base of the axon (grey) at the time of axon initiation relative to the cell centroid at 0,0 for dorsal and transverse view for ascending (n = 8 cells), descending (n = 8 cells) and peripheral axons (n = 7 cells). Lines connect centrosome and nascent axon from the same cell. R, rostral; C, caudal; A, apical; B, basal; D, dorsal; V, ventral.

Figure 3 - figure supplement 2

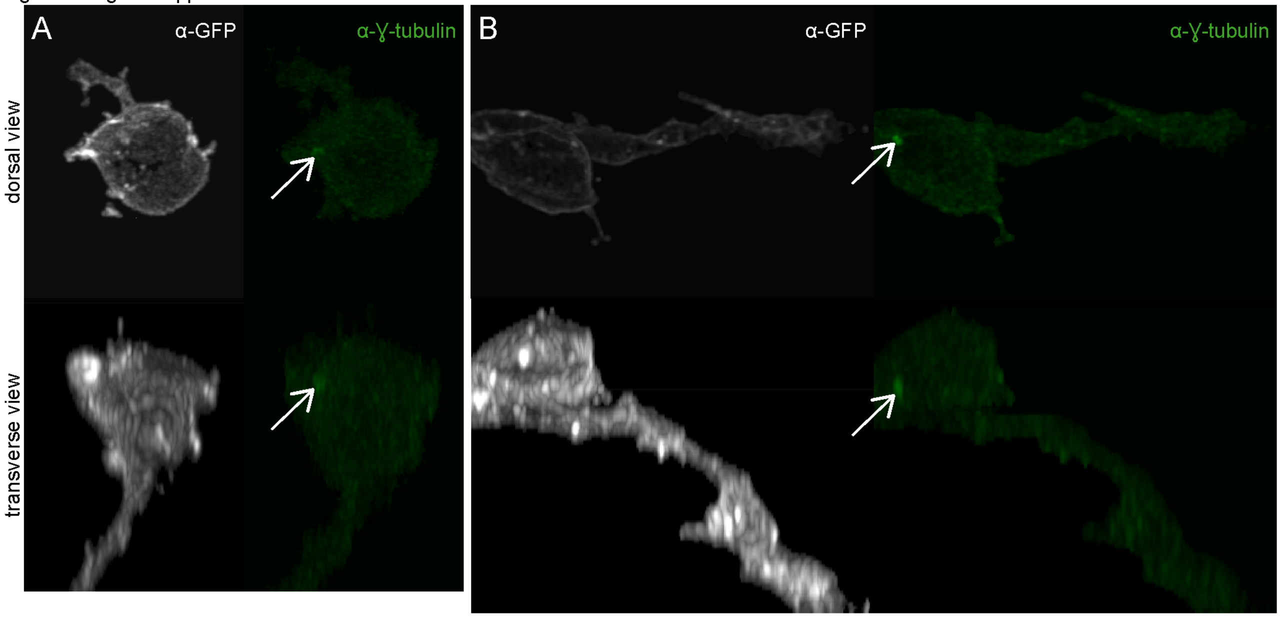

**Figure 3 – Figure supplement 2.**
(**A,B**) Two examples of neurons labelled with membrane label EGFP-CAAX in embryos processed for immunohistochemistry against GFP (grey) and ɣ-tubulin (green). ɣ-tubulin accumulations appear to correspond with the centrosome (arrows). Dorsal views are maximum projections from confocal z-stacks. Transverse views are reconstructions from confocal z-stacks.

Figure 4 - figure supplement 1

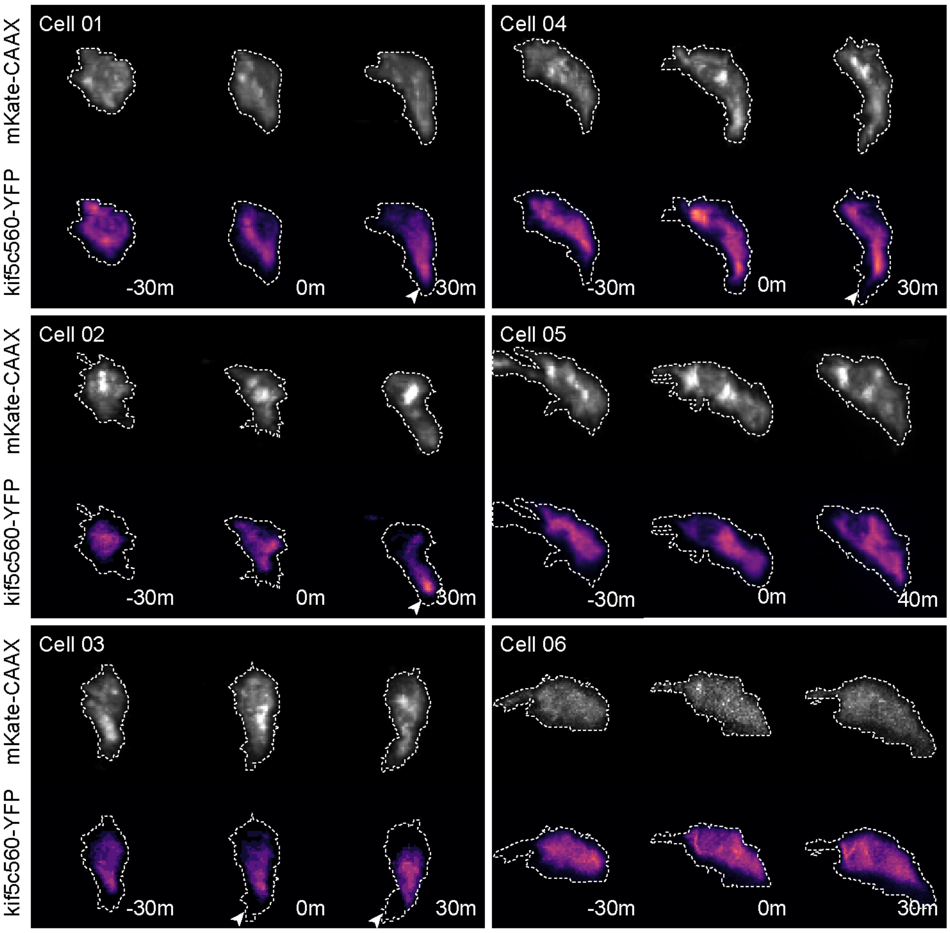

**Figure 4 – Figure Supplement 1.**
Six different neurons labelled with a membrane marker (grey) and kif5c560-YFP (magma LUT) 30 minutes before, during (0m) and 30 minutes after nascent axon initiation. Dotted line shows cell outline. Images are maximum projections of transverse reslices of confocal z-stacks.

Figure 4 - figure supplement 2

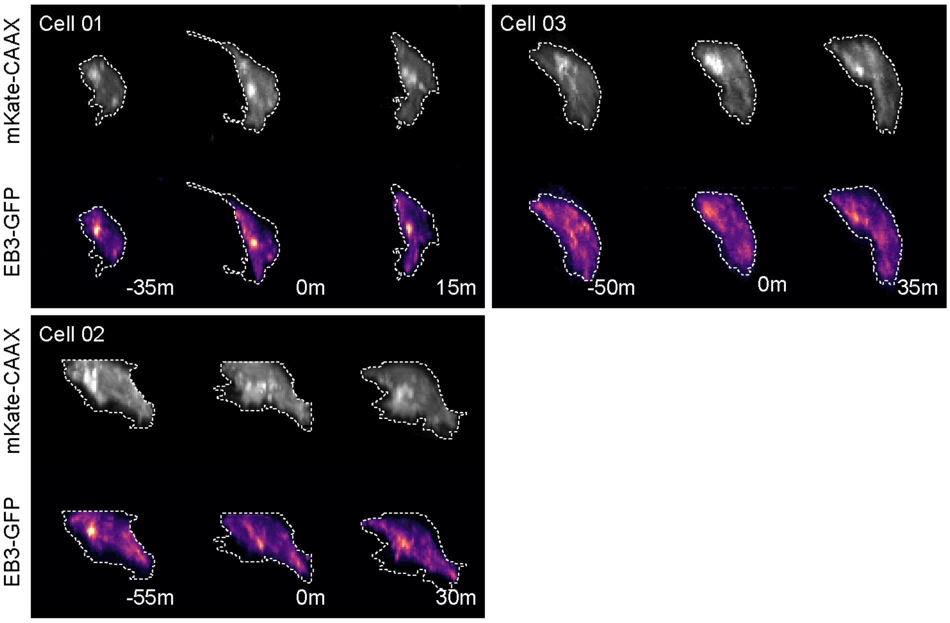

**Figure 4 – Figure Supplement 2.**
Three different neurons labelled with a membrane marker (grey) and EB3-GFP (magma LUT) 30 minutes before, during (0m) and 30 minutes after nascent axon initiation. Dotted line shows cell outline. Images are maximum projections of transverse reslices of confocal z-stacks.

Figure 4 - figure supplement 3

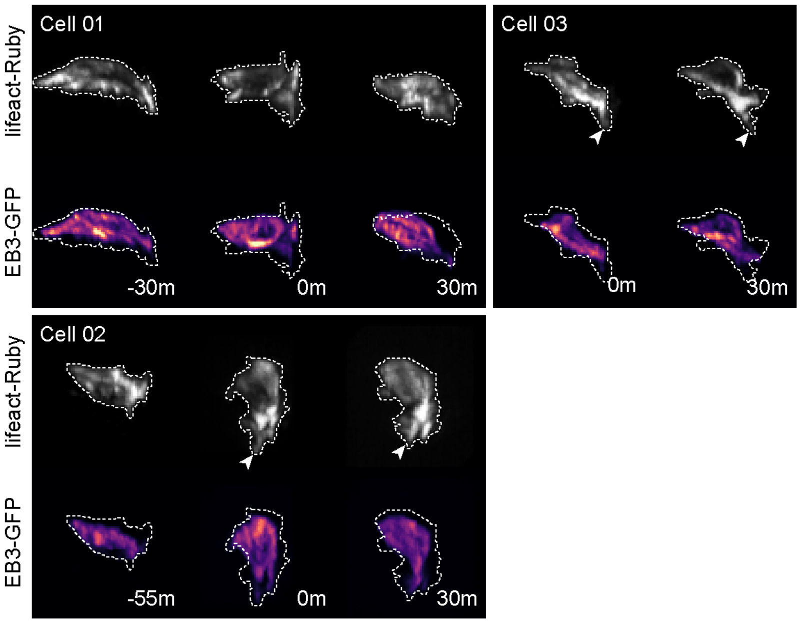

**Figure 4 – Figure Supplement 3.**
Three different neurons labelled with lifeact-Ruby (grey) and EB3-GFP (magma LUT) 30 minutes before, during (0m) and 30 minutes after nascent axon initiation. Dotted line shows cell outline. Images are maximum projections of transverse reslices of confocal z-stacks.

Figure 4 - figure supplement 4

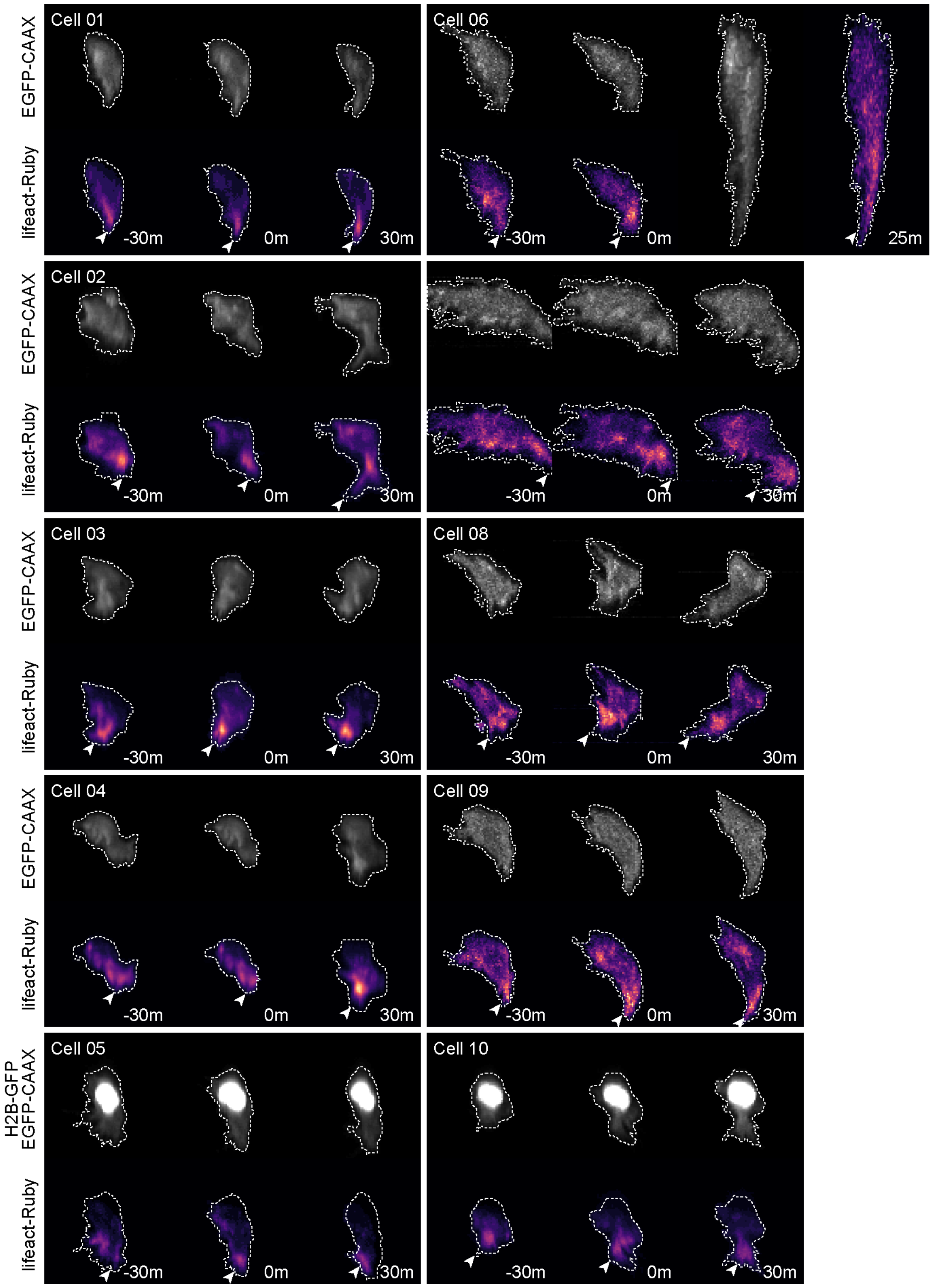

**Figure 4 – Figure Supplement 4.**
Ten different neurons labelled with a membrane marker (grey) and lifeact-Ruby (magma LUT) 30 minutes before, during (0m) and 30 minutes after nascent axon initiation. Dotted line shows cell outline. Images are maximum projections of transverse reslices of confocal z-stacks.

Figure 4 - figure supplement 5

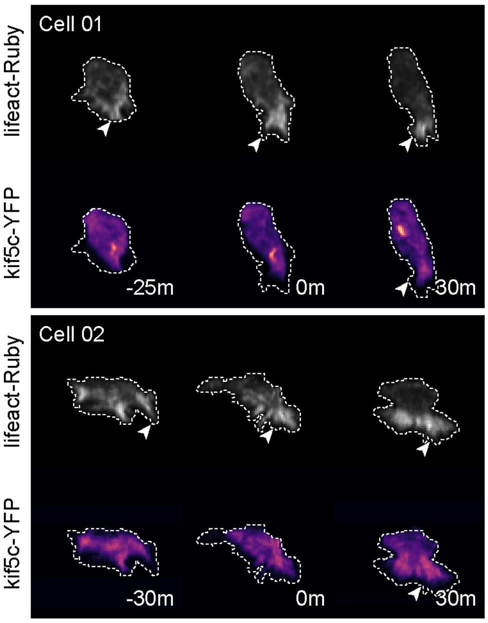

**Figure 4 – Figure Supplement 5.**
Two different neurons labelled with lifeact-Ruby (grey) and kif5c560-YFP (magma LUT) 30 minutes before, during (0m) and 30 minutes after nascent axon initiation. Dotted line shows cell outline. Images are maximum projections of transverse reslices of confocal z-stacks.

Figure 4 - figure supplement 6

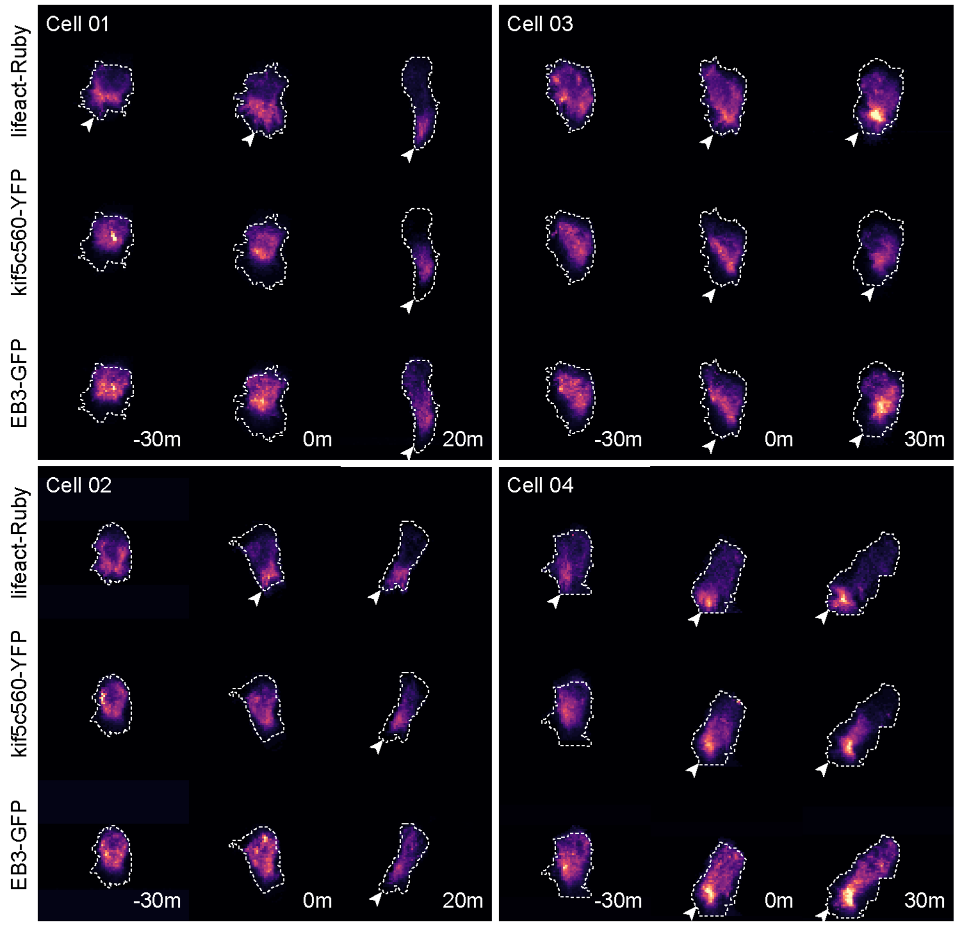

**Figure 4 – Figure Supplement 6.**
Four different neurons labelled with lifeact-Ruby, kif5c560-YFP and EB3-GFP 30 minutes before, during (0m) and 30 minutes after nascent axon initiation. Dotted line shows cell outline. Images are maximum projections of transverse reslices of confocal z-stacks.

Figure 5 - figure supplement 1

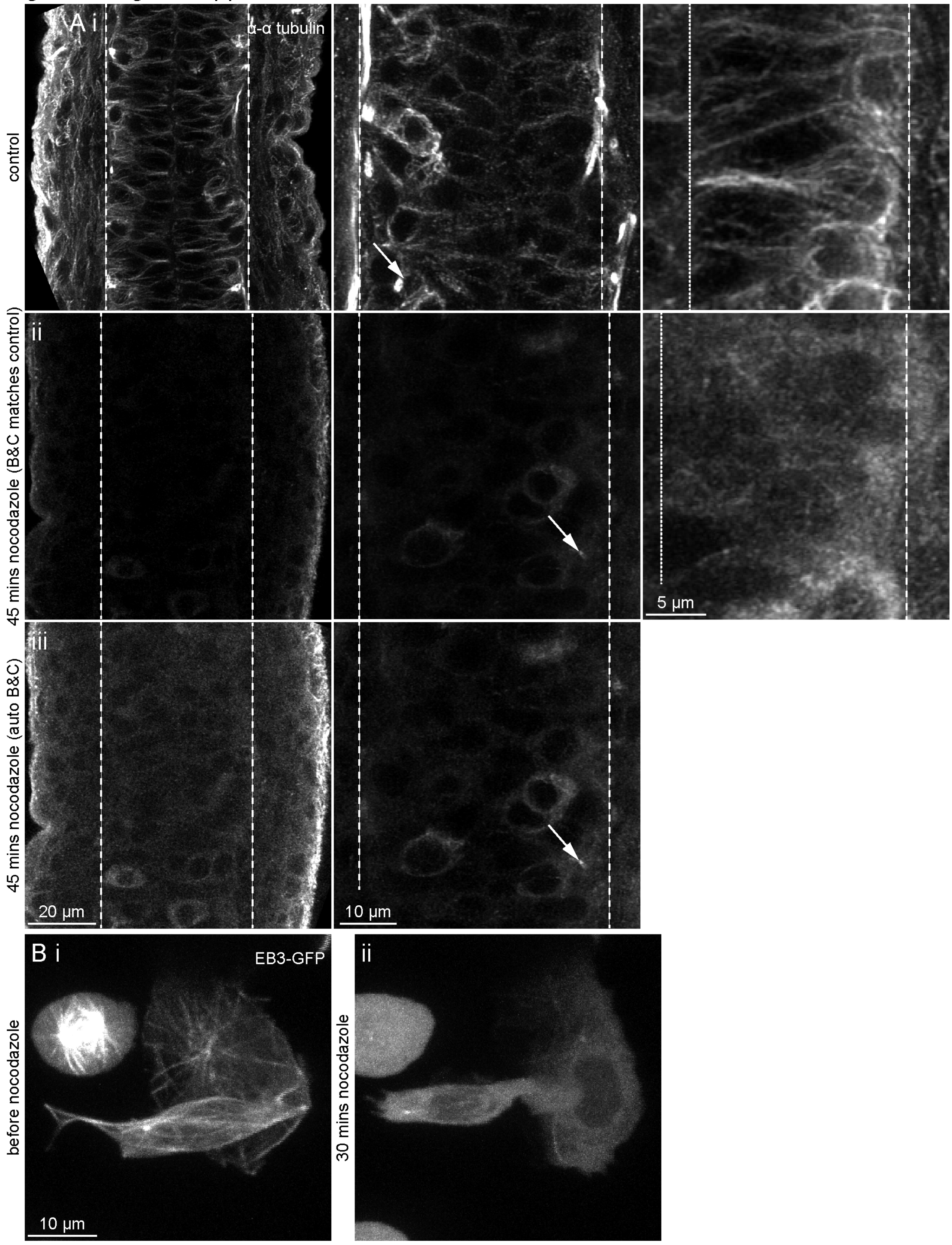

**Figure 5 – Figure Supplement 1.**

(**A**) Projections of short z-stacks showing dorsal view of zebrafish embryos incubated (**i**) in fish water (control), or (**ii, iii**) in 5 μg/mL nocodazole for 45 minutes and processed for immunohistochemistry against $\alpha$-tubulin. Three panels showing three different embryos imaged at different magnifications are shown for each condition. (**ii**) and (**iii**) show the same image but with brightness and contrast adjusted to be the same as control (ii) or automatically adjusted (iii). Control images show clear microtubule arrays in all cells. After 45 minutes of nocodazole treatment, the microtubule array is completely disrupted in neuroepithelial cells and newborn neurons, although remnants can be seen in mature neurons and their axons (arrows). Dashed lines show basal surface of neural tube; dotted lines show apical surface. Arrows = $\alpha$-tubulin labelling in established axons. (**B**) Cells labelled with EB3-GFP before and after nocodazole treatment. EB3 comets labelling microtubule plus-ends present before nocodazole treatment are not visible within 30 minutes of nocodazole treatment.

Figure 6 - figure supplement 1

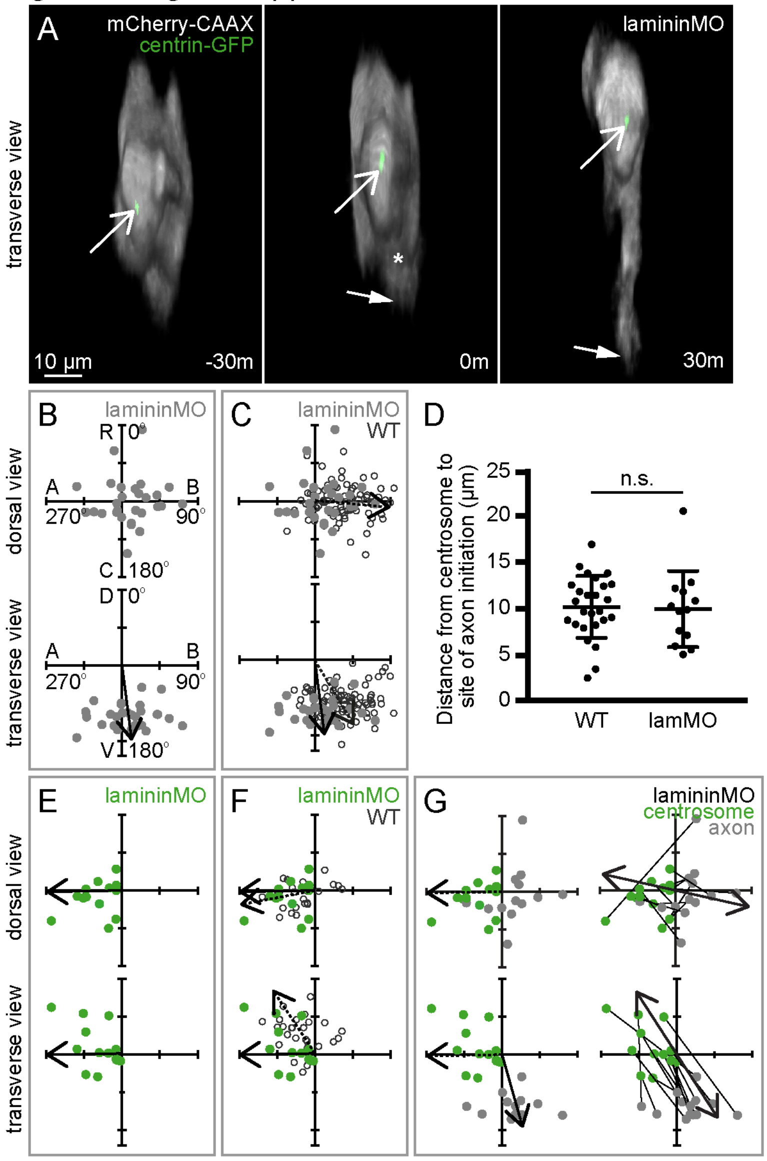

**Figure 6 – Figure Supplement 1.**
(**A**) Image sequence from confocal time lapse shows a neuron in a lamMO-injected embryo labelled with membrane (grey) and centrosome (green) markers before (-30m), during (0m) and after axon initiation (30m). Open arrows = centrosome; closed arrows = axon tip; asterisk = axon position at time of nascent axon initiation. Images are transverse reconstructions from confocal z-stacks. (**B**) Plots showing axon position on the soma relative to the cell centroid at 0,0 for dorsal and transverse view in lamMO-injected embryos (n = 29 cells). Axon position is random in dorsal view ($0.1 < P < 0.5$) but not in transverse view ($P < 0.001$). Arrow shows mean angle (transverse view mean = 172.6°; Moore's modification of Rayleigh's test). (**C**) Plots showing merge of WT (open circles) and lamMO (grey dots) axon positions on the cell body relative to cell centroid at 0,0 for dorsal and transverse views. Solid arrows show mean angle from lamMO embryos (n = 29 cells); dotted arrows show mean angles of WT embryos (n = 86 cells). Axon positions in WT and lamMO are significantly different (dorsal view $0.01 < P < 0.02$, transverse view $P < 0.001$; Batschelet's alternative to Hotelling test). (**D**) Graph showing distance between centrosome and base of axon at time of axon initiation in WT and lamMO-injected embryos. WT: n = 26 cells, mean = 10.13 µm, s.d. = 3.35. lamMO: n = 13 cells, mean = 9.9 µm, s.d. = 4.11. One-way ANOVA, $P = 0.980$. (**E**) Plots showing merge of WT (open circles) and lamMO (green dots) centrosome positions on the cell body relative to cell centroid at 0,0 for dorsal and transverse views. Solid arrows show mean angle from lamMO embryos (n = 13 cells); dotted arrows show mean angles of WT embryos (n = 26 cells). Centrosome positions are not significantly different between WT and lamMO in dorsal view ($0.2 < P < 0.5$) but are in transverse view ($0.02 < P < 0.05$; Batschelet's alternative to Hotelling test). (**F**) Plots showing the positions of the centrosome (green) and base of the axon (grey) in lamMO-injected embryos at the time of axon initiation relative to the cell centroid at 0,0 for dorsal and transverse view (n = 13 cells). Left-hand plots: centrosome position is not random (dorsal view $P < 0.001$; transverse view $P < 0.001$) and dotted arrows show mean angle of centrosome (dorsal view mean = -91.0°; transverse view mean = -91.1°); axon position is random in dorsal view ($0.05 < P < 0.1$) but not in transverse view ($P < 0.001$) and solid arrow shows mean angle of axon (transverse view mean = 164.1°; Moore's modification of Rayleigh's test). Centrosome and axon positions are significantly different (dorsal view $0.001 > P$; transverse view $0.001 > P$; Moore's test for paired data). Right-hand plots: lines connect centrosome and nascent axon from the same cell. Double-headed arrows show average slope of vectors linking centrosome and nascent axon and are not random (dorsal view $P < 0.001$, mean = 102.6°; transverse view $P < 0.001$, mean = 147.1°).

Figure 6 - figure supplement 2

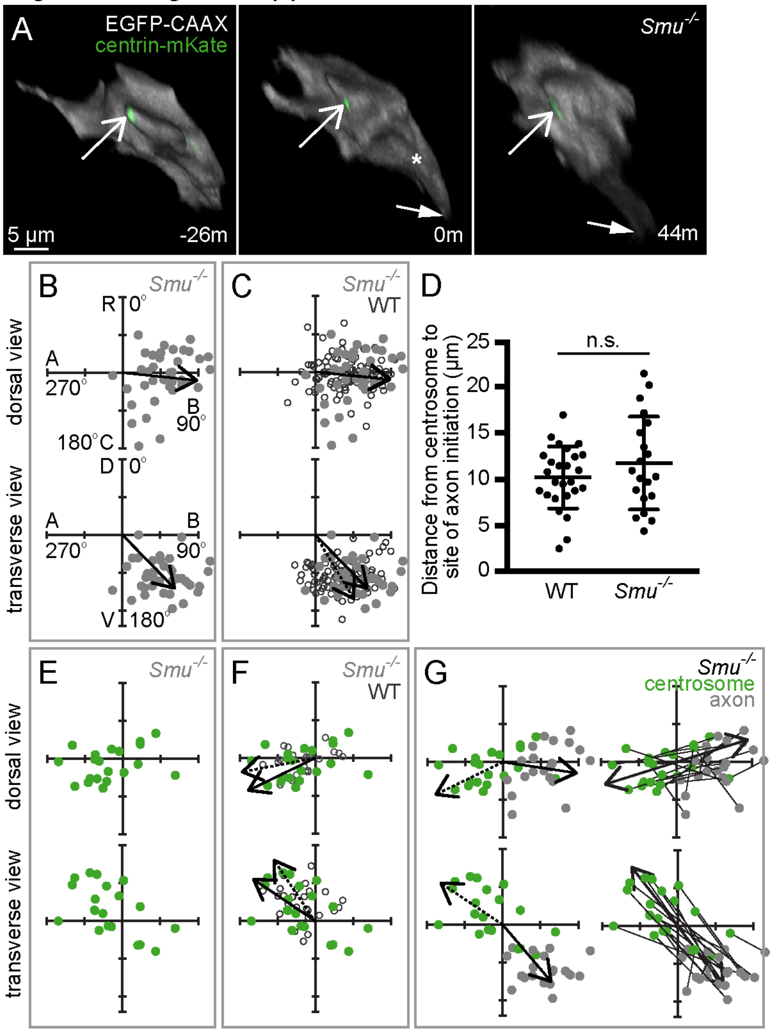

**Figure 6 – Figure Supplement 2.**
(**A**) Image sequence from confocal time lapse shows a neuron in a *Smu*$^{-/-}$ embryo labelled with membrane (grey) and centrosome (green) markers before (-26m), during (0m) and after axon initiation (44m). Open arrows = centrosome; closed arrows = axon tip; asterisk = axon position at time of nascent axon initiation. Images are transverse reconstructions from confocal z-stacks. (**B**) Plots showing axon position on the soma relative to the cell centroid at 0,0 for dorsal and transverse view (n = 38 cells) in *Smu*$^{-/-}$ embryos. Axon position is not random (dorsal view 0.001 > *P*; transverse view *P* < 0.001). Arrows show mean angle (dorsal view mean = 98.24°; transverse view mean = 135.7°; Moore's modification Rayleigh's test). (**C**) Plots showing merge of WT (open circles) and *Smu*$^{-/-}$ (grey dots) axon positions on the cell body relative to cell centroid at 0,0 for dorsal and transverse views. Solid arrows show mean angle from *Smu*$^{-/-}$ embryos (n = 38 cells) and dotted arrows show mean angles of WT embryos (n = 86 cells). Although axon positions in WT and *Smu*$^{-/-}$ are significantly different in dorsal view (0.02 < *P* < 0.05), this is not the case in transverse view (0.05 < *P* < 0.1; Batschelet's alternative to Hotelling test). (**D**) Graph showing distance between centrosome and base of axon at time of axon initiation in WT and *Smu*$^{-/-}$ embryos. WT: n = 26 cells, mean = 10.13 μm, s.d. = 3.348. *Smu*$^{-/-}$: n = 20 cells, mean = 11.68 μm, s.d. = 5.027. Student's two-tailed test, *P* = 0.216. (**E**) Plots showing merge of WT (open circles) and *Smu*$^{-/-}$ (green dots) centrosome positions on the cell body relative to cell centroid at 0,0 for dorsal and transverse views. Solid arrows show mean angle from *Smu*$^{-/-}$ embryos (n = 20 cells) and dotted arrows show mean angles of WT embryos (n = 26 cells). Centrosome positions are not significantly different between WT and *Smu*$^{-/-}$ embryos (dorsal view 0.5 < *P*; transverse view 0.2 < *P* < 0.5; Batschelet's alternative to Hotelling test). (**F**) Plots showing the positions of the centrosome (green) and base of the axon (grey) in *Smu*$^{-/-}$ embryos at the time of axon initiation relative to the cell centroid at 0,0 for dorsal and transverse view (n = 20 cells). Left-hand plots: centrosome position is not random (dorsal view 0.025 < *P* < 0.05; transverse view 0.01 < *P* < 0.025) and dotted arrows show mean angle of centrosome (dorsal view mean = -115.8°; transverse view mean = -56.1°); axon position is not random (dorsal view *P* < 0.001; transverse view *P* < 0.001) and solid arrows show mean angle of axon (dorsal view mean = 98.5°; transverse view mean = 139.5°). Centrosome and axon positions are significantly different (dorsal view 0.001 > *P*; transverse view 0.001 > *P*; Moore's test for paired data). Right-hand plots: lines connect centrosome and nascent axon from the same cell. Double-headed arrows show average slope of vectors linking centrosome and nascent axon, which are not random (dorsal view 0.001 < *P* < 0.005, mean = 72.7°; transverse view *P* < 0.001, mean = 142°; Moore's modification of Rayleigh's test).
