## Supplementary material for "Actin-based protrusions lead microtubules during stereotyped axon initiation in spinal neurons *in vivo*": Video legends

**Video 01. Early steps of differentiation in the zebrafish spinal cord.** Maximum projection of confocal timelapse, dorsal view. A neuron labelled with a membrane marker extends two transient protrusions along the basal surface (open arrows). They are retracted, along with the apical attachment (arrowhead), before the axon is extended (closed arrow).

**Video 02. Axon initiation is highly stereotyped.** Transverse reconstruction of confocal time lapse. The neuron, labelled with a membrane marker, first extends multiple transient pre-axonal protrusions (arrowheads). It then extends a nascent axon (0 mins), which is maintained for approximately 30 minutes before axon growth begins. Arrows show axon tip.

**Video 03. Axon initiation can be separated from axon growth.** 3D reconstructions of confocal time lapse in transverse and lateral views. A DoLA neuron has stereotypical baso-ventral axon initiation. After nascent axon formation, the axon turns and grows rostrally to establish its characteristic axon trajectory. Arrows show axon tip.

**Video 04. Most zebrafish spinal neurons do not undergo apical abscission.** Maximum projection of confocal time lapse, dorsal view. A neuron (asterisk) labelled with a membrane marker retracts its apical process from the apical surface of the spinal cord. No abscission is observed. Solid line shows position of apical surface; arrowheads show tip of retracting apical process.

**Video 05. The centrosome and cilium stay in close proximity and move together towards the basal surface.** Maximum projection of confocal time lapse, dorsal view. Cilia are tagged with GFP and centrosomes are shown in magenta. Arrowheads show one cilium-centrosome pair as it moves from the apical surface towards the basal surface.

**Video 06. The centrosome is not close to the nascent axon during axon initiation.** Transverse reconstruction of confocal time lapse. A neuron is labelled with membrane (grey) and centrosome (green) markers. Open arrows show centrosome position before, during (0 mins) and after axon initiation. Closed arrows show axon tip; asterisk indicates base of axon.

**Video 07. Axon marker Kif5c560 is enriched in the nascent axon and growth cone.** Transverse reconstruction from confocal time lapse. A neuron is labelled with a membrane marker (grey) and Kif5c560-YFP (yellow). Kif5c560 is localised throughout the cell body before nascent axon initiation, then is gradually enriched in the nascent axon and then growth cone during axon growth. Arrows show axon tip; arrowheads indicate kif5c accumulation.

**Video 08. Microtubule plus-end marker EB3 is not enriched in the nascent axon.** Transverse reconstruction from confocal time lapse. A neuron is labelled with lifeact-Ruby to mark F-actin (greys) and EB3-GFP to mark microtubule plus-ends (green). Most EB3 is located in the cell body before axon initiation and during (0 mins) nascent axon establishment. EB3 is first enriched in the axonal growth cone during axon growth (arrowheads). Arrows show axon tip.

**Video 09. Few microtubule plus-ends enter the nascent axon.** Maximum projection from confocal time lapse, dorsal view. A neuron is labelled with EB3-GFP to mark microtubule plus-ends. Each field shows the same neuron at different timepoints. Frames are every 5 seconds. Compared to the amount of microtubules in the cell body, few microtubule plus-ends grow into the basal-most section of the cell before axon initiation (-30 mins) or into the nascent (0 mins, 30 mins), but do reach the tip of the growing axon (60 mins). Arrows show axon tip.

**Video 10. F-actin is persistently localised baso-ventrally before nascent axon initiation.** Transverse reconstruction from confocal time lapse. A neuron is labelled with a membrane marker (grey) and lifeact-Ruby to mark F-actin (green). Frames are every 2 minutes. Actin is persistently localised baso-ventrally before a persistent nascent axon protrusion. Arrowheads at -60 mins show persistent actin localisation, arrows at 0 mins show axon tip.

**Video 11. An actin-rich nascent axon-like protrusion can develop during nocodazole treatment.** Transverse reconstruction from confocal time lapse. A neuron is labelled with a membrane marker (grey) and lifeact-Ruby to mark F-actin (green). The neuron has not yet extended an axon before nocodazole treatment (-10 mins). All protrusions are retracted upon application of nocodazole (0 mins). During nocodazole treatment the neuron extends multiple small, transient non-axonal protrusions in many directions as well as a longer, persistent, actin-rich nascent axon-like protrusion ventrally (from 130 mins).

**Video 12. Laminin provides a basal cue for axon initiation.** Transverse reconstruction of confocal time lapse. A neuron is labelled with membrane (grey) and centrosome (green) markers. The axon is extended ventrally while the centrosome is away from the site of axon initiation (0 mins). Asterisk shows position of axon initiation; open arrows show centrosome position; closed arrows show axon tip.
